# Reduced polymerase pausing compensates for increased chromatin accessibility in the aging liver

**DOI:** 10.1101/2022.02.25.481984

**Authors:** Mihaela Bozukova, Dora Grbavac, Chrysa Nikopoulou, Peter Tessarz

## Abstract

Regulation of gene expression is tightly linked to the organization of the mammalian genome. With age, chromatin alterations occur on all levels of genome organization, accompanied by changes in the gene expression profile. However, little is known about the changes on the level of transcriptional regulation with age. Here, we used a multi-omic approach and integrated ATAC-, RNA- and NET-seq to identify age-related changes in the chromatin landscape of murine liver and to investigate how these are linked to transcriptional regulation. We provide the first systematic inventory of the connection between aging, chromatin accessibility and transcriptional regulation in a whole tissue. We observe that aging in murine liver is accompanied by an increase in chromatin accessibility at promoter regions of protein-coding genes. Yet, although promoter accessibility is a requirement for transcription, the increased accessibility does not result in enhanced transcriptional output. Instead, aging is accompanied by a decrease of promoter-proximal pausing of RNA polymerase II (Pol II). We propose that these changes in transcriptional regulation are due to a reduced stability of the pausing complex and may represent a mechanism to compensate for the age-related increase in chromatin accessibility in order to prevent aberrant transcription.

## INTRODUCTION

Epigenetic drifts as well as changes in chromatin architecture and transcriptional programs commonly occur during mammalian aging (Benayoun *et al*, 2015; Maleszewska *et al*, 2016; Benayoun *et al*, 2019) and contribute to the general decline in physiological function that is usually observed in older individuals (López-Otín *et al*, 2013). The importance of epigenetic and transcriptional alterations in the aging process is underscored by several recent studies that employed partial reprogramming – an inherently epigenetic phenomenon – to rejuvenate several tissues in a variety of different aging models. Such rejuvenation phenomena do not only positively impact physiology, but also established a more youthful epigenetic state as measured by changes in the DNA methylation pattern (Lu *et al*, 2020; Chondronasiou *et al*, 2022), the levels of histone modifications (Ocampo *et al*, 2016) and the transcriptome itself (Lu *et al*, 2020). In summary, epigenetic and transcriptional states change with age and are directly linked to physiological fitness. Importantly, however, all studies addressing aging conducted up to now have investigated transcriptional changes using either microarrays or RNA-seq. These technologies measure steady-state levels of mRNA, which depend on the rate of nascent transcription and mRNA degradation (Nikopoulou *et al*, 2019) – both of which might change upon aging. Recent advances have made it possible to directly monitor nascent transcripts, using approaches such as GRO-seq (Core *et al*, 2008) and NET-seq (Mayer *et al*, 2015), but so far, these technologies have been only used in tissue culture.

Local chromatin architecture is an important regulator of transcription (Venkatesh & Workman, 2015). Nucleosomes create physical barriers to RNA polymerase (Pol II) and transcription factors (Petesch & Lis, 2012). Thus, for initiation of transcription, the promoter region needs to be rendered accessible, which is achieved by the combined action of pioneering transcription factors (TFs) (Zaret & Carroll, 2011) and chromatin remodelers (Venkatesh & Workman, 2015). The size of the nucleosome-free region positively correlates with TF binding and subsequent gene activation (Scruggs *et al*, 2015). During transcription initiation, Pol II is recruited to the accessible promoter region through the concerted action of general transcription factors, resulting in the sequential formation of the preinitiation complex (PIC), before phosphorylation of Ser5 and Ser7 of the C-terminal domain (CTD) of Pol II triggers promoter escape (Wong *et al*, 2014). However, Pol II stalls after the initial transcription of about 20-60 nucleotides, in a process termed promoter-proximal pausing. Over the last decade, genome-wide studies have demonstrated that Pol II pausing is a ubiquitous step in the transcription cycle of Drosophila and mammalian genes (Muse *et al*, 2007; Zeitlinger *et al*, 2007), which serves as an additional regulatory layer. Paused polymerase is thought to allow further finetuning of transcription and the integration of various upstream signals. Indeed, ES cells that have decreased pausing efficiency became refractory to differentiation cues, highlighting the role of pausing in regulating signaling cascades in response to differentiation cues (Williams *et al*, 2015). In addition, paused Pol II is important to maintain nucleosome-depleted regions around promoters (Core & Adelman, 2019). Stable promoter-proximal Pol II pausing is facilitated by binding of the pause-inducing factors DSIF (DRB sensitivity-inducing factor), a heterodimer consisting of SPT4 and SPT5 as well as NELF (negative elongation factor) to Pol II (Wada *et al*, 1998; Wu *et al*, 2003; Lee *et al*, 2008). To release paused Pol II into productive elongation, CDK9 as part of the of P-TEFb complex (Marshall & Price, 1995), phosphorylates NELF, the DSIF subunit SPT5 as well as Ser2 of the CTD of Pol II. SPT5 phosphorylation transforms DSIF from an inhibitory to a stimulating elongation factor (Bernecky *et al*, 2017) and triggers the dissociation of NELF and progression of Pol II into productive elongation.

Here, we analyzed how local chromatin structure and transcription initiation are altered in the aging liver using ATAC-seq together with a NET-seq protocol adapted for tissues and in combination with publicly available RNA-and ChIP-seq data. We demonstrate that aging in the liver is characterized by a strong increase in promoter accessibility. However, this increase in accessibility is not reflected in an increased transcriptional output on nascent or steady-state level. Interestingly, the overlap of differentially regulated steady-state and nascent RNA is minor, indicating that post-transcriptional processes play a major role in shaping the transcriptome during the aging process. Despite the overall increase in promoter accessibility, we show that the levels of promoter-proximal pausing globally decline with age. Mapping of SPT4 binding to chromatin revealed that DSIF is recruited less efficiently to the promoter-proximal region in aged livers. Furthermore, several lines of evidence suggest that transcriptional initiation is unaltered with age. Together, these data indicate a general loss of stability of the Pol II pausing complex upon aging, which we propose might mitigate the effects of the overall increase in promoter accessibility to maintain a relatively young-like transcription state in the aging liver.

## RESULTS

### Promoter regions are more accessible with age

To understand how genome accessibility changes on a local scale with age, we performed ATAC-seq using liver tissue from independent biological replicates of each young (3-month-old) and aged (18-month-old) mice (Figure 1A). The generated ATAC-seq datasets were of high quality and displayed the characteristic fragment size distribution (Supplementary Figure 1A-D). As a first-level analysis, we performed principal component analysis (PCA). PCA revealed that samples from the same age group (young or aged) formed two distinct clusters mostly distinguishable by the first two principal components, which explain more than 70 % of the variance in the data (Supplementary Figure 1E). Differential accessibility analysis revealed that only 8.43 % of analyzed regions (4,691 out of 55,669) were differentially accessible with age (Figure 1B).

**Figure 1:**
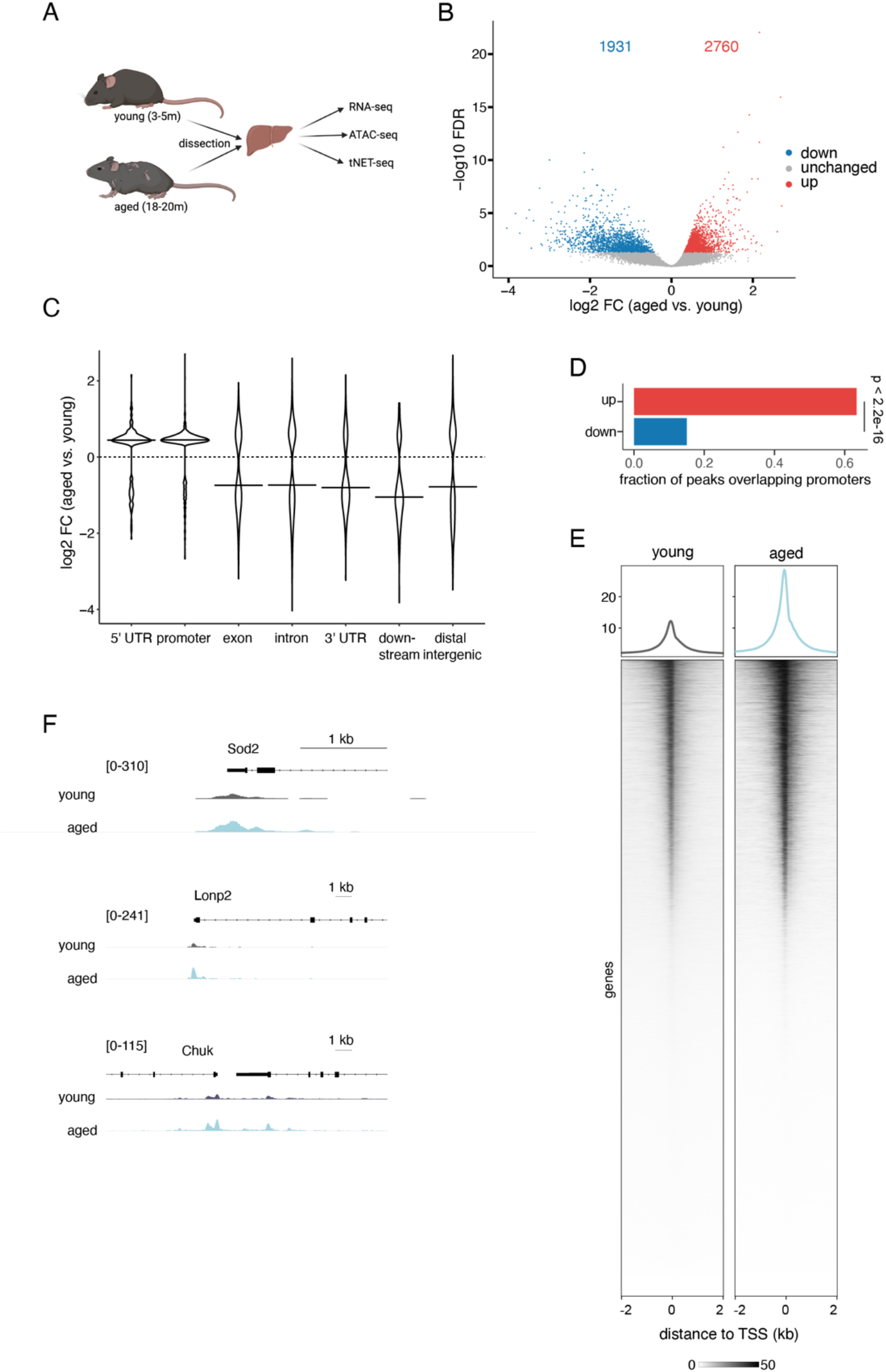
Chromatin accessibility at promoter regions increases with age in murine liver tissue. A. Schematic of experimental set-up. B. Volcano plot of differentially accessible genomic regions comparing liver tissue of aged relative to young mice (FDR < 0.05, Wald test). Increased accessibility: 2,760 regions (red), decreased accessibility: 1,931 regions (blue), regions with unchanged accessibility: 50,978 (grey). C. Genomic locations of differentially accessible sites. Y-axis represents the log2-fold change (log2 FC) in accessibility in the liver of aged versus young animals. D. Proportion of differentially accessible sites overlapping promoters (defined as TSS ± 200 bp). P-value was calculated using a two-sided, two proportion z-test. E. Heatmap and average intensity profiles of promoter accessibility over all annotated TSSs in the mouse genome. Read densities of merged biological replicates were normalized to 1x coverage. F. Representative genome browser views of promoters with increased accessibility in aged mice. Peak intensity range is indicated in brackets. Reads from biological replicates were merged and normalized to 1x coverage.

Of these, 2,760 sites became more accessible, while 1,931 sites showed decreased accessibility with age. This slight trend towards an overall increased accessibility prompted us to further zoom into the differentially accessible sites by investigating their genomic location. We observed that sites with increased accessibility in the aged liver were on average closer to annotated TSSs compared to less accessible sites, suggesting an age-related increase in the accessibility of promoter regions (Supplementary Figure 2A). Indeed, promoter regions and 5’ UTRs became more accessible with age, while the accessibility of distal intergenic and genic sites decreased (Figure 1C). In fact, the majority of regions (63.4 %), for which we detected an increased accessibility, were located within annotated promoter regions (Figure 1D). In contrast, only 15.0 % of regions with decreased accessibility in aged animals overlapped with known promoter regions (Figure 1D). This age-related increase in chromatin accessibility at promoter regions was detectable both on a metagene (Figure 1E) and single-gene level (Figure 1F). Furthermore, gene ontology (GO) enrichment analysis revealed that genes with more accessible promoters in aged liver tissue were involved in metabolic processes such as amino acid metabolism (Supplementary Figure 2B, Supplementary Table 1). In contrast, genes with decreased promoter accessibility with age were linked to nucleosome assembly and organization (Supplementary Figure 2C, Supplementary Table 2). Together, our results indicate that liver aging is accompanied by an increased chromatin accessibility at promoter regions.

### Aging has a modest effect on the transcriptional output

Given the central role of promoter accessibility in transcription, we next asked whether the observed age-related increase in promoter accessibility affects the transcriptional output. For this, we assessed steady-state transcription using publicly available liver RNA-seq data from the Tabula Muris Senis Consortium (Schaum *et al*, 2020). For a more detailed look into the temporal dynamics of gene expression changes with age, we exploited the availability of gene expression data from multiple age groups and included, in addition to young and old (3-and 18-months of age, respectively), also middle-aged (12-month-old) mice (Figure 1A). Gene expression patterns in aged liver were clearly distinct from those of young and middle-aged animals as observed by PCA (Supplementary Figure 3A). Differential expression analysis revealed that the expression of the majority of genes did not significantly change with age (Supplementary Figure 3B, C). Thus, gene expression in the liver appears to be relatively resistant to aging, consistent with observations from single-cell RNA-seq data of liver hepatocytes from the Tabula Muris Consortium (Tabula Muris Consortium, 2020).

To directly assess the effect of promoter accessibility on the transcriptional output, we integrated our ATAC-seq with the RNA-seq data. Interestingly, the age-related increase in promoter accessibility is not directly reflected in increased transcriptional output (Figure 2A). Of note, standard RNA-seq measures steady-state mRNA levels, which are determined by both synthesis and degradation rates. To exclude the contribution of mRNA degradation, we next focused exclusively on nascent transcription.

**Figure 2:**
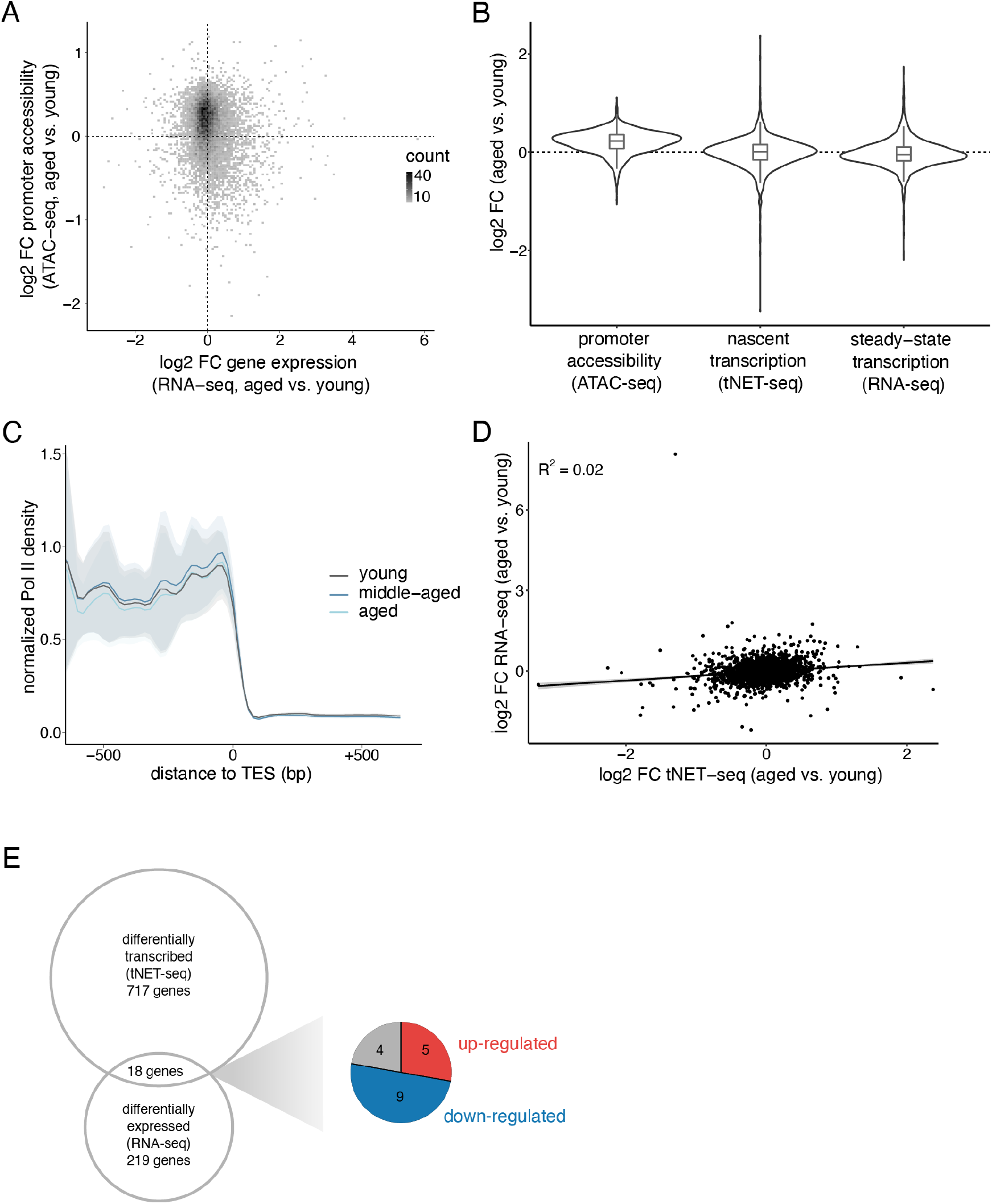
Aging has a subtle effect on transcriptional output in murine liver tissue. A. Scatter plot of the log_2_ fold change in gene expression (RNA-seq) and promoter accessibility (ATAC-seq) in aged versus young mice. Promoter region defined as TSS +/- 200 bp. B. Violin and boxplots of changes in promoter accessibility (ATAC-seq), nascent transcription (gene-body Pol II density, tNET-seq) and steady-state transcription (RNA-seq) in aged versus young mice. Only genes present in all three datasets are included here (n = 2,693 genes). Promoter defined as TSS +/- 200 bp. C. Metagene profile showing normalized Pol II densities (tNET-seq) at TESs. Read densities were normalized to 1x coverage. Solid lines represent mean values and shading indicates the 95 % confidence interval. D. Correlation between changes in nascent (gene-body Pol II density, tNET-seq) and steady-state transcription (RNA-seq) of aged versus young animals. E. Venn diagram of differentially transcribed (gene-body Pol II density, tNET-seq) and differentially expressed genes (RNA-seq) in aged versus young mice. Only significantly changed genes (FDR < 0.05, Wald test) were included. Up-or down-regulated genes in both datasets are indicated in red and blue, respectively. Genes exhibiting divergent changes are indicated in grey.

To investigate age-induced changes in nascent transcription on a genome-wide level, we employed native elongating transcript sequencing (NET-seq) (Mayer *et al*, 2015). NET-seq quantitatively maps the position of transcriptionally engaged Pol II with single-nucleotide resolution and strand specificity. Importantly, NET-seq does not require previous labeling of mRNA and thus, is ideally-suited for measuring nascent transcription in tissues. For application in murine liver tissue, we modified the protocol to include efficient tissue homogenization and nuclei isolation in one single step. We named this new, modified version of the original protocol tissue NET-seq (tNET-seq). We performed tNET-seq using freshly isolated liver tissue from three independent biological replicates from young (3-month-old), middle-aged (12-month-old) and aged (18-month-old) mice. These age groups match those of the RNA-seq data, enabling direct integration of the two datasets. tNET-seq libraries showed high reproducibility among biological replicates (Supplementary Figure 4A, B), indicating the high robustness of the approach.

Using tNET-seq, we assessed how nascent transcription is affected by age. For this, we stringently defined a set of protein-coding genes that do not overlap with other transcription units within 2.5 kb of the TSS and TES and are longer than 2 kb (n = 12,460 genes). We further included only genes with sufficient coverage (RPKM > 1, n = 3,280 genes). Using these genes, we assessed Pol II density at TES as a proxy for nascent transcriptional output. Consistent with the modest changes we observed in steady-state expression levels, the nascent transcriptional output was also only mildly affected with age (Figure 2B, C). Yet, there were clear differences between the nascent transcriptome profiles of young and aged liver, assessed by PCA (Supplementary Figure 4C). To further dissect these age-related changes in nascent transcription, we quantified the nascent transcript levels by assessing Pol II density within gene bodies. Compared to young animals, we found 11 % (367 out of 3,278) and 22 % (717 out of 3,278) of genes to be differentially transcribed in middle-aged and aged liver, respectively (Supplementary Figure 4D, E). There was an equal distribution between up-and down-regulated nascent transcripts, highlighting the fact that there is no global unidirectional shift (i.e., increase or decrease) in the nascent transcription with age. Overall, these results demonstrate that while promoter accessibility is a requirement for transcription, its increased accessibility in the aging liver does not automatically result in elevated transcriptional output.

### Age-related changes in nascent transcription are not directly mirrored in steady-state mRNA levels

To link nascent with steady-state transcription, we directly compared tNET-seq and RNA-seq data sets. The moderate positive correlation between the normalized signal indicated an overall agreement between the two data sets (Supplementary Figure 5). However, we observed only a weak positive correlation between age-related changes in nascent and steady-state transcription (Figure 2D). In fact, only 18 genes were found to be both differentially transcribed (tNET-seq) and differentially expressed (RNA-seq) with age (Figure 2E). Together, these results demonstrate that age-related changes in nascent transcription do not necessarily reflect the steady-state mRNA levels and highlight the important contribution of post-transcriptional regulatory processes, such as mRNA degradation, to steady-state mRNA levels.

### Age-related changes in nascent transcription lack a clear signature

Considering the limited number of genes convergently changing on nascent and steady-state level, we next focused our attention exclusively on the nascent transcription and investigated whether differentially transcribed genes exhibit a specific signature in their gene features. We found no correlation between age-related changes in nascent transcription and gene length (Supplementary Figure 6A), exon length (Supplementary Figure 6B) or exon number per gene (Supplementary Figure 6C).

To dissect temporal trajectories of changes in nascent transcription, we performed differential testing using a likelihood ratio test. Considering young, middle-aged and aged animals, we found 29 % of genes (953 out of 3,278) to be differentially transcribed. Trajectory analysis of these differentially transcribed genes resulted in four distinct gene clusters with similar patterns and distinct functional enrichment (Figure 3): Genes in cluster 1 showed an age-related increase in nascent transcription and were linked to mRNA processing and splicing (Figure 3A). Genes in clusters 2 and 3 exhibited an overall decrease and increase, respectively, and were linked to metabolic processes. While cluster 2 was functionally linked to xenobiotic and carboxylic acid catabolic processes, genes in cluster 3 were involved in lipid, carbohydrate or glutathione metabolic processes. This highlights the diversity of age-related, metabolic changes in the highly metabolic liver tissue. The smallest gene cluster (cluster 4) contained very few genes (Figure 3D). Interestingly, some of these genes (*Parp14, Irgm1, Nlrc5*) are linked to the interferon gamma response, which is known to be involved in the aging of the liver and is connected to the development of “inflammaging” (Singh *et al*, 2011).

**Figure 3:**
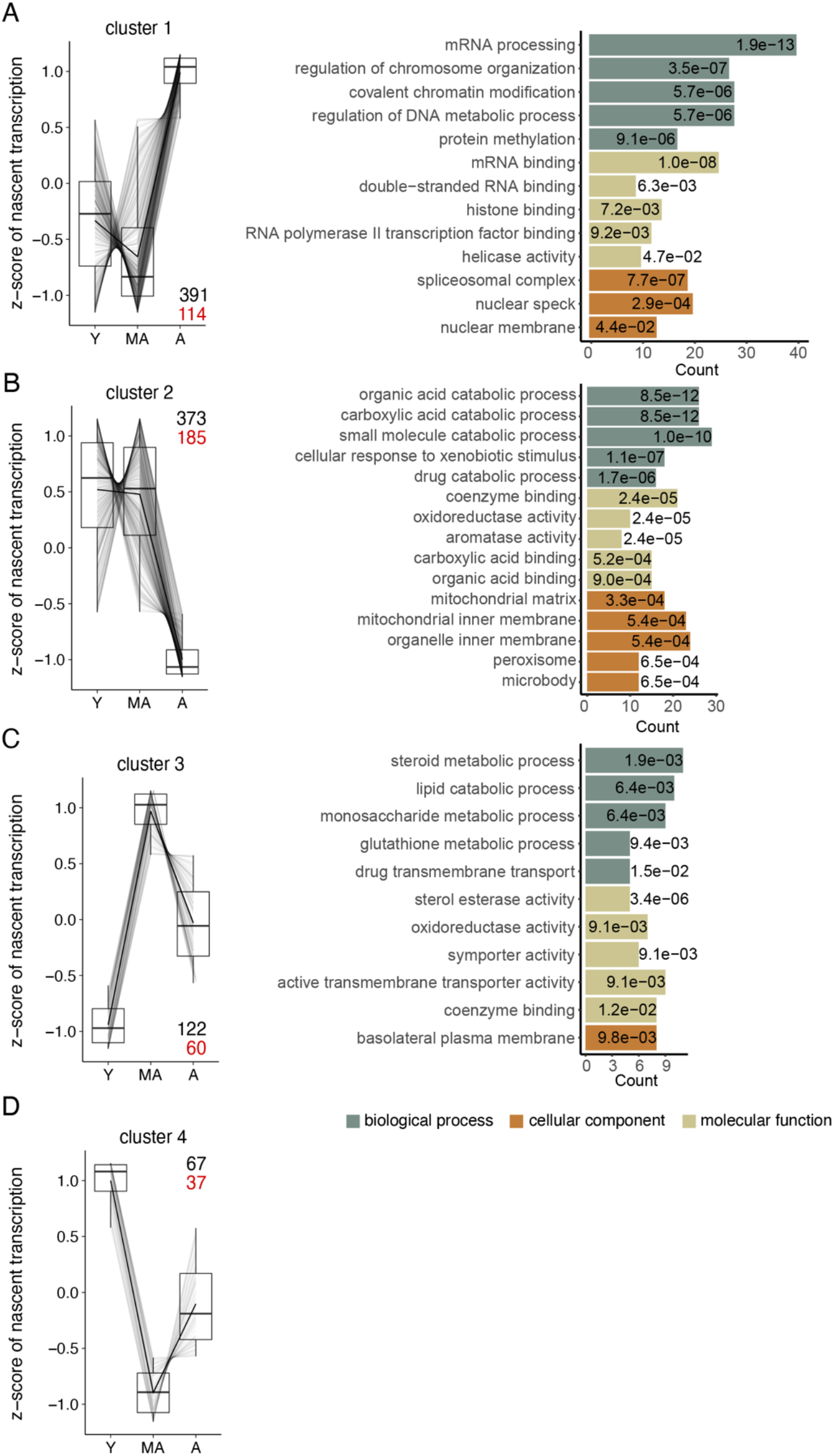
Trajectory analysis of differentially transcribed genes in aged versus young mice (n = 953 genes, likelihood ratio test, DESeq2) resulted in four distinct gene clusters. In each cluster, the total number of genes and the number of “aging genes” are designated in black and red, respectively. Nascent transcript levels are reported in the parallel coordinate plot on the left as z-scores of gene-body Pol II density. GO term analysis for each gene cluster is depicted on the right. Only the top five enriched GO terms ranked by adjusted p-value (Benjamini-Hochberg (BH) procedure, FDR < 0.05) are displayed. BH-adjusted p-values are reported for each GO term inside the respective bar. Y, young; MA, middle-aged; A, aged.

To understand the importance of these differentially transcribed genes in the context of murine aging, we cross-referenced them to a recently published resource of “global aging genes” based on single-cell RNA-seq data from the Tabula Muris Senis Consortium (Zhang *et al*, 2021). A large fraction of the hepatocyte-specific aging genes (45 %, 1,158 out of 2,569) was present in our tNET-seq dataset that we used for differential transcription testing and trajectory analysis. We found 34 % of these aging genes (396 out of 1,158) to be differentially transcribed with age, which is in the same order of magnitude as the overall differentially transcribed genes (29 %, 953 out of 3,278). The differentially transcribed “aging genes” were not enriched in any particular cluster suggesting no common pattern in the changes of their nascent transcription (Figure 3, Supplementary Tables 3-6). Taken together, these analyses reveal distinct age-related trajectories in the nascent transcriptome with a large fraction of the previously defined “aging genes” being altered.

Yet, the changes in nascent transcription were not congruent and unidirectional, suggesting that nascent transcription alone does not control the coordinated global aging behavior at tissue and organismal level reported earlier. This corroborates the modest correlation between tNET-seq and RNA-seq we observed (Figure 2D), implying that the other processes, such as mRNA processing and degradation might shape the coordinated gene expression signature detected within and across tissues.

### Promoter-proximal Pol II pausing decreased with age

The discrepancy between the increased promoter accessibility, the overall modest change in transcriptional output with age and the lack of a clear signature in terms of common functionalities of differentially transcribed genes, led us to focus on the initial stages of transcription at the promoter region, which is where much of the transcriptional regulation occurs. (t)NET-seq is a powerful approach for investigating promoter-proximal Pol II pausing, which is a major regulatory step of transcription. To quantify promoter-proximal pausing, we calculated the pausing index (PI), which is the ratio of the average Pol II density in the promoter region (defined here as TSS +/-200 bp) to that in the gene body (defined here as TSS + 200 bp to TES -200 bp). The PI provides a measure of the magnitude of promoter-proximal Pol II density relative to that in the gene body. Based on the PI values in young animals, we classified genes into three pausing categories with high (PI ≥ 3), moderate (1.5 < PI < 3) or low (PI ≤ 1.5) pausing. 69.7 % of the analyzed genes were either highly or moderately paused in young animals (Supplementary Figure 7A). This corroborates promoter-proximal pausing as a wide-spread phenomenon in the liver, consistent with previous reports in other organisms and cell types. To investigate alterations in promoter-proximal Pol II pausing with age, we generated metagene profiles of a 1-kb region around the genes’ TSSs (Figure 4A). Notably, we observed a progressive age-related decrease in promoter-proximal Pol II pausing (Figure 4A). This decrease, however, was not accompanied by changes in Pol II density in the immediate 5’ gene bodies (Figure 4A) or at the TES as a proxy for transcriptional output (Figure 2C). The age-related decrease in promoter-proximal pausing is also reflected in a progressive decrease in PI (Figure 4B). Interestingly, highly paused genes (PI > 3 in young mice) experienced the highest decrease in promoter-proximal Pol II pausing with age (Supplementary Figure 7B).

**Figure 4:**
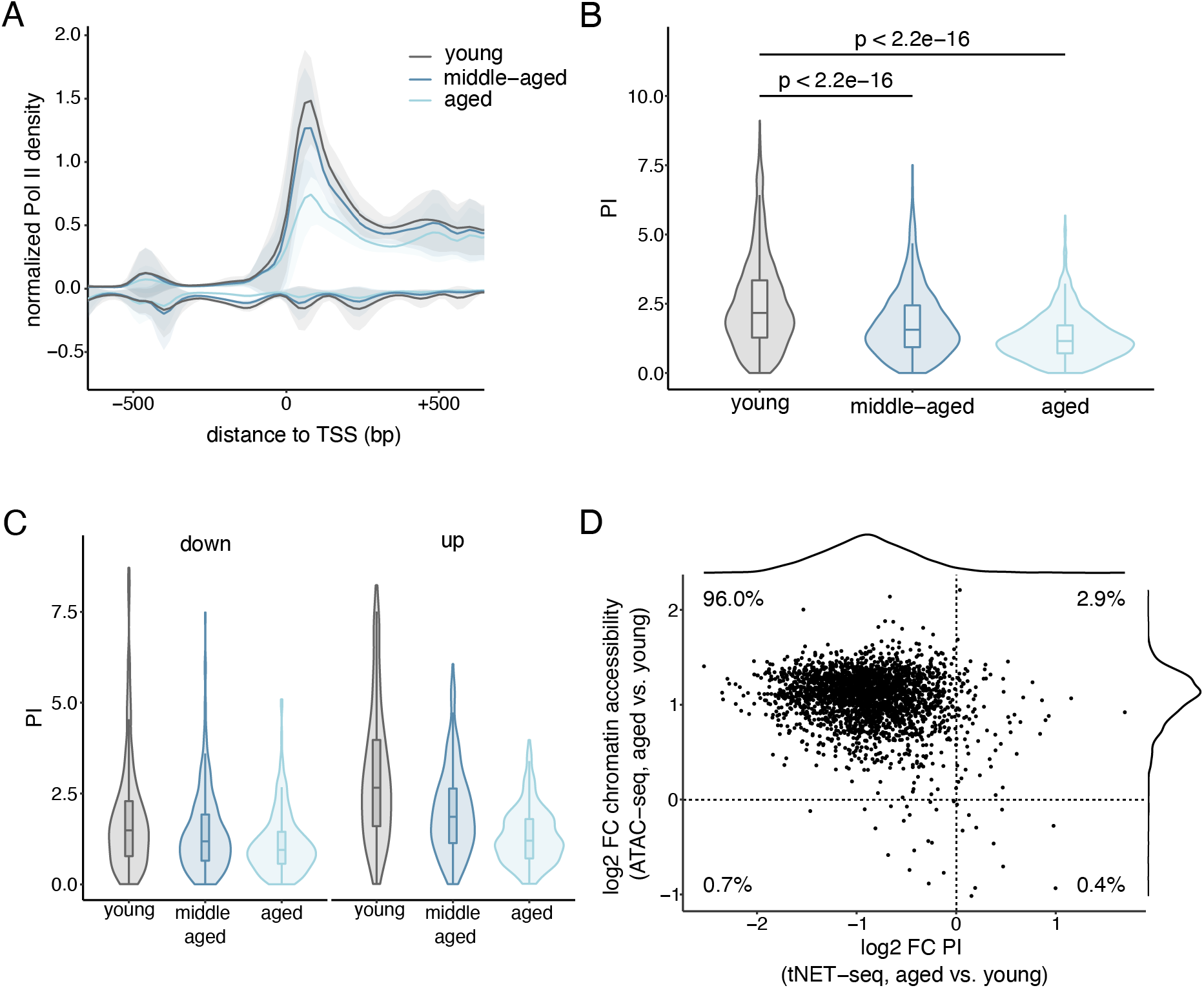
Promoter-proximal Pol II pausing decreases with age in murine liver. A. Metagene profile of normalized Pol II densities (tNET-seq) at the promoter-proximal region of all genes included in the analysis. Read densities were normalized to 1x coverage. Solid lines represent mean values and shading indicates the 95 % confidence interval. B. Violin and box plots of PI values in young, middle-aged and aged animals. p-values were calculated using a two-sided Wilcoxon rank-sum test. C. Violin and box plots showing the PI in young, middle-aged and aged animals. Only genes found to be differentially transcribed in aged versus young mice are included and grouped by the direction of change (down-regulated, left panel; up-regulated, right panel). D. Scatter plot of the change in PI (tNET-seq) and promoter accessibility (ATAC-seq) in aged versus young mice. Promoter region defined as TSS +/- 200 bp.

Changes in the PI depend on both changes in the numerator (promoter-proximal Pol II) and denominator (gene-body Pol II). Therefore, we considered the possibility that the age-related decrease in PI may be related to the levels of nascent transcription. Hence, we grouped the gene set into equal-sized groups based on their level of nascent transcription in young mice (Supplementary Figure 7C). The fold changes in PI between all groups were similar, indicating that the age-related decrease in PI occurs irrespective of the gene’s expression level. This was further confirmed by the observation that differentially transcribed genes consistently exhibited a lower PI in aged animals (Figure 4C). This is true even for the down-regulated genes, corroborating that the observed effect was specific to the promoter-proximal region (numerator of PI) and could not be explained solely by changes in nascent transcription (denominator of PI). Overall, these findings reveal an age-related decrease in promoter-proximal Pol II pausing affecting the majority of the analyzed genes.

The local chromatin landscape at promoter regions directly modulates transcriptional regulation and progression of Pol II by regulating the accessibility of the region. To investigate the relationship between promoter accessibility and promoter-proximal pausing, we integrated the ATAC-seq and tNET-seq datasets. The majority of investigated promoters (96.0 %) exhibited an increased chromatin accessibility with a concomitant decrease in promoter-proximal Pol II pausing (Figure 4D). In light of the modest age-related changes in transcriptional output (Figure 2B), this points towards a compensatory mechanism, in which alterations at the step of promoter-proximal pausing might counteract the increased accessibility of the promoter regions, thereby maintaining transcriptional fidelity.

### Aging affects the stability of the Pol II pausing complex

How can the age-related decrease in promoter-proximal Pol II pausing be explained? Focusing solely on the transcription process, we can envision three possible scenarios: A decrease in promoter-proximal Pol II pausing can be a consequence of a decreased initiation rate, a decreased duration of pausing, or a combination of both.

Alterations in transcription initiation would be detected as decreased promoter-proximal Pol II pausing. (t)NET-seq does not allow for directly quantifying initiating Pol II, since the nascent RNA needs to be at least 20 bp long for adapter ligation and unambiguous read alignment to the genome. Therefore, we focused on enhancer regions instead, which regulate transcription initiation levels at multiple levels (Beagrie & Pombo, 2016). For identifying active enhancers in murine liver, we integrated our ATAC-seq data with publicly available histone ChIP-seq data from liver tissue of young (3 month), middle-aged (12 months) and aged (29 months) mice (Benayoun *et al*, 2019). This resulted in the identification of 8,855 high-confidence, putative enhancers, that were present in all three age groups. The majority of these enhancers are located in intronic regions (Supplementary Figure 8A), consistent with a recent report highlighting the predominantly intronic location of tissue-specific enhancers (Borsari *et al*, 2021). Having identified these liver-specific enhancers, we then assessed enhancer activity using chromatin accessibility as a proxy. We observed an increased enhancer accessibility in aged compared to young animals (Figure 5A). In fact, 21.3 % (587 out of 2,760) of the genomic regions that were more accessible with age were located within enhancer regions (Figure 5B). In contrast, only 6.7 % (129 out of 1,931) of sites that exhibited a significant decrease in accessibility with age were found within enhancer regions (Figure 5B).

**Figure 5:**
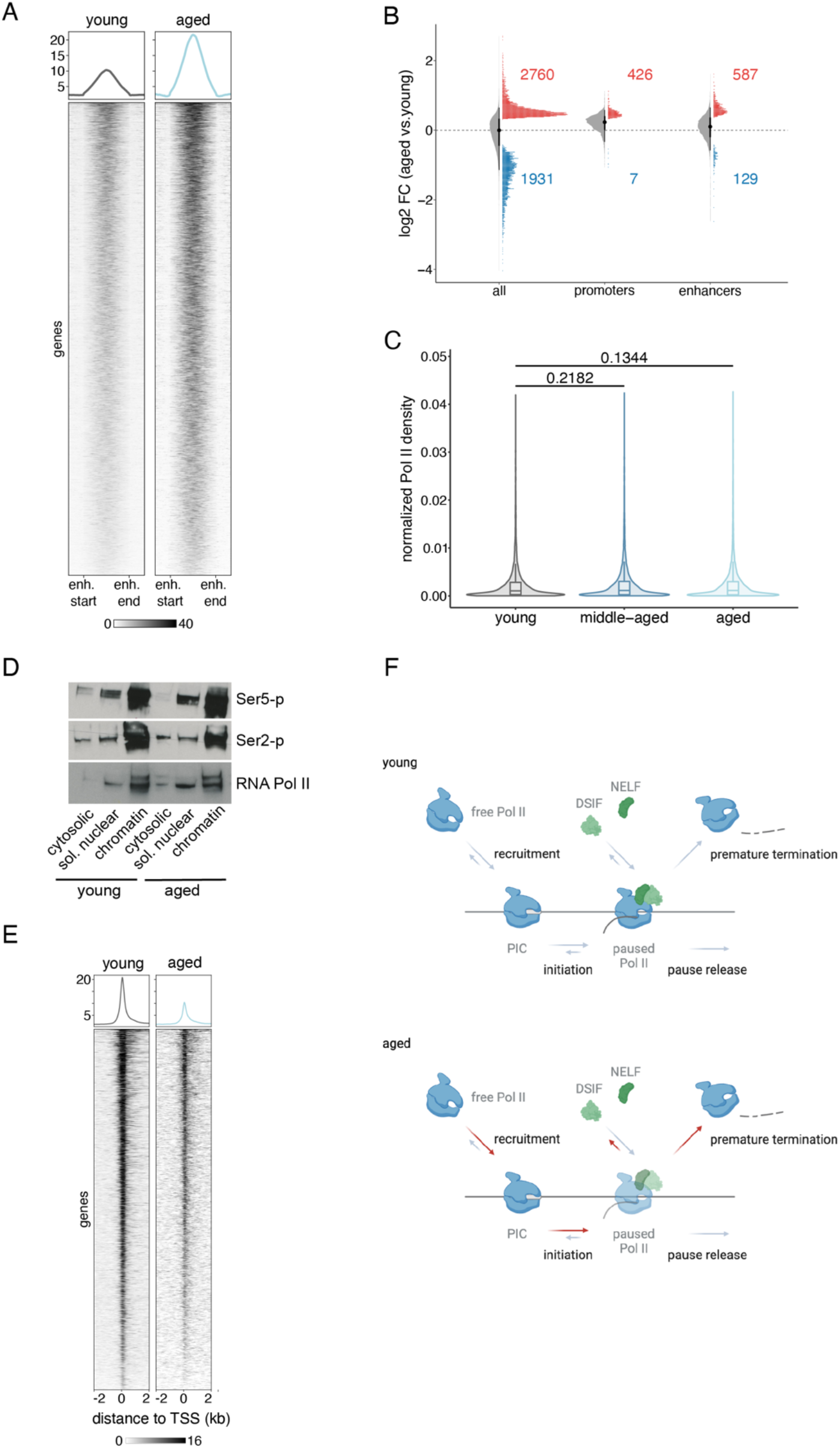
Aging affects the stability of the Pol II pausing complex rather than transcription initiation. A. Heatmap and intensity profiles of promoter accessibility over active enhancers in liver (n = 8,855). B. Raincloud plots of changes in accessibility in aged versus young liver assessed by ATAC-seq. Raincloud plots are hybrid plots. Here, the violin and boxplots visualize the log2-fold change for all accessible genomic sites (“all”) or accessible sites overlapping tNET promoters or enhancers. The dot plot highlights regions significantly changed with age in each category (FDR < 5 %). The numbers in the plot denote significantly up- or down-regulated regions in each category. C. Violin and boxplots of Pol II density at enhancer regions as a means of quantifying eRNA production. p-values were calculated using a two-sided Wilcoxon rank-sum test. D. Western Blots of cellular fractionations of young and aged liver samples probed with the indicated antibodies. E. Heatmaps and average intensity profiles of SPT4 ChIP-seq signal at TSSs of tNET genes in young and aged animals. Read densities were normalized to 1x coverage. F. Proposed model for the age-related changes that occur in the liver. Increased chromatin accessibility and indirect assessment of initiation suggest that this step is not affected, or rather increased as indicated by the red arrows. On the other hand, decreased pausing in the promoter-proximal region and lower recruitment of SPT4 suggest that the pausing complex is less stable upon aging, resulting in premature termination of transcription (indicated by red arrow).

This trend of more enhancer sites becoming more accessible with age compared to those with decreased accessibility mirrors the accessibility changes at promoters (Figure 1E). As a second proxy for enhancer activity, we quantified enhancer RNA transcription, since eRNA production and enhancer activity have been reported to highly correlate on a genome-wide scale. (t)NET-seq captures nascent RNAs, including short-lived RNA species like eRNAs, which are not detected with sufficient coverage by RNA-seq. We quantified eRNA production by assessing Pol II levels at the identified enhancer regions. We observed no age-related change in eRNA production (Figure 5C, Supplementary Figure 8B). Overall, our data point towards unaltered or increased, but not decreased initiation rates. Finally, we assessed the levels of Ser5 and Ser2 phosphorylation on the CTD of Pol II. As decreased promoter proximal pausing occurs globally, an overall decline in initiation should be visible at the level of CTD phosphorylation. In line with our analysis on enhancer activity, we did not observe any change in the level of Ser5 nor Ser2 phosphorylation (Figure 5D). Taken together, these data in combination with the overall increase in promoter accessibility strongly suggest that transcription initiation is not compromised in aged livers but is likely enhanced.

The second possible explanation for the age-related decrease in promoter-proximal Pol II pausing is an age-related decrease in Pol II dwell time. We first assessed the expression levels of important transcriptional regulators involved in the early stages of transcription. We observed no change in the expression of these factors with age (Supplementary Figure 8C). However, while the expression of these transcription regulators was not affected by age, their recruitment to chromatin might have been altered. To assess this, we performed ChIP-seq for SPT4, a component of the pause-inducing factor DSIF. We observed a decreased occupancy of SPT4 at TSSs of tNET genes in aged animals (Figure 5E). Thus, while their expression does not seem to be affected with age, the recruitment and binding of important pausing factors to chromatin might be altered, resulting in changes in promoter-proximal Pol II pausing.

This is consistent with the decreased level of promoter-proximal Pol II pausing we observed using tNET-seq (Figure 4A, B). These results suggest that the age-related decrease in promoter-proximal Pol II pausing might be explained by an altered stability of the pausing complex in aged animals. The dissociation of Pol II from chromatin at this early stage of transcription might antagonize the effect of the age-related increase in promoter accessibility and thus, a potential increase in Pol II recruitment and transcription initiation. Overall, a potential increase in initiation and a concomitant decrease in stability of the pausing complex would lead to an unchanged transcriptional output (Figure 5F).

## DISCUSSION

The rapid development of omics technologies in recent years have significantly advanced our understanding of the epigenome and its role in transcriptional regulation. Using freshly isolated liver tissue from young, middle-aged and aged mice, we provide a comprehensive analysis of age-related changes in the local chromatin landscape and the nascent transcriptome. The combination of multiple genome-wide sequencing techniques (tNET-seq, ATAC-seq, RNA-seq, ChIP-seq) creates a framework for investigating the connection between chromatin and transcriptional regulation upon ageing.

In recent years, several studies have interrogated age-related changes in the chromatin landscape of different tissues and cell systems. Chromatin accessibility analysis of aged CD8+ T-cells from human donors revealed a decrease in promoter accessibility (Moskowitz *et al*, 2017; Ucar *et al*, 2017). Recently, an overall more compacted chromatin architecture has also been observed in aged murine neutrophils (Lu *et al*, 2021) and mesenchymal stem cells (Pouikli *et al*, 2021). Contrasting these observations of decreased chromatin accessibility, a study reports no major changes in accessibility in aged murine B precursor cells (Koohy *et al*, 2018). Thus, it is likely that age-related alterations in chromatin accessibility are highly tissue- and cell-type specific, which may, at least in part, explain these contrasting observations. We find that in murine liver tissue, a specific fraction of the genome undergoes age-related changes. Particularly promoter regions gain accessibility. Importantly, we do not observe changes in the expression of histone genes, indicating that the increase in promoter accessibility is not simply a consequence of decreased nucleosomes. In support of this, no global changes in nucleosome occupancy (assessed using H3 occupancy as a proxy) have been reported in aged murine liver tissue; only at a subset of loci either increased or decreased occupancy was observed (Chen *et al*, 2020). This is also consistent with another report, in which MNase-seq of aged mouse liver revealed no global changes in nucleosome occupancy (Bochkis *et al*, 2014).

Despite the increase in promoter accessibility, we observe only modest effects on both the nascent and steady-state transcriptome with age. This highlights that the expression levels of most genes are generally preserved in aged liver, consistent with the notion that liver tissue seems to be more refractory to aging (Zhang *et al*, 2021) as assessed by an aging score based on scRNA-seq data from the Tabula Muris Consortium (Tabula Muris Consortium, 2020). Hepatocytes from different age groups exhibited similar aging scores, suggesting that their transcriptional programs are marginally affected by age (Zhang *et al*, 2021). This is in contrast, for instance, to immune and stem cells that exhibited a stronger increase in aging scores, reflecting the higher turnover rates of these cell types. Intriguingly, we observed only a small overlap between differentially expressed and differentially transcribed genes upon aging, with more widespread changes identified in the nascent transcriptome. A similar discordance has been observed in total versus nuclear RNA-seq analyses of young and aged murine B precursor cells (Koohy *et al*, 2018). Thus, changes in the nascent transcriptome might be buffered through post-transcriptional mechanisms affecting RNA stability. This is consistent with recent results analyzing single-cell RNA-and ATAC-seq data of aged murine liver, which also highlighted the important contribution of post-transcriptional processes (Nikopoulou *et al*, 2021).

Despite the modest age-related effects on transcriptional output, we observed a strong decrease in promoter-proximal Pol II pausing. How can this age-related decrease in promoter-proximal Pol II pausing be explained? We provide data demonstrating reduced recruitment of the DSIF subunit SPT4 to the promoter-proximal region, suggesting that the pausing complex is less stable in aged hepatocytes. However, we cannot distinguish between a reduced recruitment to or an increased dissociation from chromatin. How DSIF is recruited to Pol II remains enigmatic. A recent mass spectrometry analysis revealed that the proto-oncogene MYC recruits the second DSIF subunit, SPT5, to promoters, thereby promoting the assembly of SPT5 with Pol II and controlling the processivity of Pol II elongation complexes (Baluapuri *et al*, 2019). Furthermore, MYC also regulates pause release by recruiting the P-TEFb subunit CDK9 to promoters (Rahl *et al*, 2010). Considering the link of MYC to inflammation and cancer (Greten & Grivennikov, 2019), it would be interesting to further explore its role in the context of aging and promoter-proximal Pol II pausing. Besides an altered recruitment of the pausing factors, aging might affect the dissociation of the pausing complex from chromatin. SPT5 depletion using RNAi in Drosophila S2 cells led to a loss of promoter-proximal Pol II, without a release into productive elongation, suggesting increased levels of promoter-proximal transcription termination (Henriques *et al*, 2018). This mirrors effects observed in NELF-depleted cells, where turnover of promoter-proximal Pol II was more rapid (Gilchrist *et al*, 2010; Henriques *et al*, 2013; Shao & Zeitlinger, 2017). Thus, DSIF and NELF are central to regulating the stability of paused Pol II. Recent studies using rapid inducible protein depletion have provided an even more fine-grained view of the roles of DSIF and NELF. Acute SPT5 depletion led to destabilization and degradation of promoter-proximal Pol II (Aoi *et al*, 2021; Hu *et al*, 2021). In contrast, acute NELF depletion triggered premature termination of promoter-proximally paused Pol II (Aoi *et al*, 2020). Both of these distinct pathways result in removal of Pol II from chromatin. These lines of evidence suggest that our results could be explained by a reduced stability of paused Pol II with age. It is currently actively debated on how much of the paused Pol II complex proceeds into productive elongation versus promoter-proximal termination (Core & Adelman, 2019). Our results suggest an age-related shift of the balance towards promoter-proximal termination. Future work will elucidate the magnitude with which these contribute to the observed age-related decrease in promoter-proximal Pol II pausing. Of particular interest is the Integrator complex, whose RNA endonuclease activity has been implicated in promoter-proximal transcription termination (Skaar *et al*, 2015; Elrod *et al*, 2019). Notably, both NELF and DSIF can associate with the Integrator complex (Yamamoto *et al*, 2014).

Overall, to explain the age-related decrease in promoter-proximal Pol II pausing, we propose the following model (Figure 5F): With age, chromatin accessibility at promoters of protein-coding genes increases. This increase in chromatin accessibility might lead to an initial increase in transcription initiation and thus transcriptional output. Such increase in transcriptional output, when passing a threshold, might then trigger a negative feedback loop impinging on the stability of the pausing complex to prevent further aberrant transcription and maintain transcription fidelity.

## MATERIALS AND METHODS

### Mouse husbandry

Animals were bred and housed in the mouse facility of the Max Planck Institute for Biology of Ageing. Experimental procedures were approved by the State Office North Rhine-Westphalia, Germany (LANUV) and are in accordance with institutional and national guidelines. Young (3-month-old), middle-aged (12-month-old) and aged (18-month-old) male C57BL/6N mice were used. Mice were provided with *ad libitum* standard rodent diet and water.

### ATAC-seq library preparation

ATAC-seq libraries were prepared following the Omni-ATAC protocol (Corces *et al*, 2017) using liver tissue of three young and aged biological replicates. Nuclei were isolated from 50-100 mg of liver tissue. For this, samples were incubated with 500 μl of of nuclei isolation buffer I (494 μl of nuclei EZ lysis buffer, 55 μl of 10x DNAseI buffer and 1 μl of DNAse I) for 15 min on ice. Then, 250 μl of nuclei EZ lysis buffer (Sigma-Aldrich, NUC101) were added and samples were vortexed vigorously (4 cycles of 2 seconds “on” and 1 second “off”). After centrifuging the samples at 500 x g for 5 min, the pellets were incubated with 250 μl of nuclei isolation buffer II (246.5 μl of nuclei EZ lysis buffer, 27.5 μl of 10x DNAseI buffer and 1 μl of DNAse I) for 10 min on ice. Then, 500 μl of nuclei EZ lysis buffer (Sigma-Aldrich, NUC101) were added and samples were vortexed vigorously as described above. After centrifuging the samples at 500 x g for 5 min, the pellets were incubated in 500 μl of nuclei EZ lysis buffer (Sigma-Aldrich, NUC101) for 20 min on ice. Residual debris was removed by filtration through a 40-μm cell strainer followed by centrifugation at 500 x g for 5 min. After washing the nuclei pellet with 1x PBS, nuclei were counted using a hemocytometer. 100,000 nuclei were then used for ATAC-seq library preparation following the Omni-ATAC protocol. Libraries were sequenced in paired-end mode on an Illumina HiSeq 2500 platform.

### ChIP-seq library preparation

ChIP-seq libraries were prepared from two independent biological replicates per age group (young and aged). Unless otherwise stated, the entire procedure was performed at 4°C or on ice. Freshly harvested liver tissue was washed four times with ice-cold 1x PBS (Gibco), cut on ice into small pieces and washed three more times with 1x PBS. The tissue was then crosslinked with 1 % formaldehyde and homogenized in a pre-chilled Dounce tissue homogenizer using a loose pestle (15 strokes). After incubation for 10 min rocking at room temperature, the crosslinking reaction was quenched with the addition of glycine to a final concentration of 0.125 M. After incubation for 5 minutes rocking at room temperature, the samples were centrifuged at 3,260 x g for 5 minutes and the supernatant was discarded. 300 mg of cross-linked tissue were lysed in 2 ml lysis buffer (50 mM Hepes pH 7.9, 140 mM NaCl, 1 mM EDTA, 10 % glycerol, 0.5 % NP-40, 0.25 % Triton x-100, 0.5 µg/ml leupeptin, 0.7 µg/ml pepstatin A, 0.5 mM PMSF) and homogenized in a pre-chilled Dounce tissue homogenizer using both a loose and tight pestle (15 strokes each). After addition of 10 ml of lysis buffer, the samples were incubated on ice for 20 minutes and centrifuged at 3,260 x g for 5 minutes. Nuclei pellets were washed by resuspending twice in 10 ml wash buffer (10 mM Tris pH 8.1, 200 mM NaCl, 1mM EDTA, 0.5mM EGTA, 0.5 µg/ml leupeptin, 0.7 µg/ml pepstatin A, 0.5 mM PMSF, 5 mM sodium butyrate) and centrifuging at 3,260 x g for 5 min, and then washing with 4 ml shearing buffer (0.1 % SDS, 1 mM EDTA, 10 mM Tris pH 8.0, 0.5 µg/ml leupeptin, 0.7 µg/ml pepstatin A, 0.5 mM PMSF, 5 mM sodium butyrate) without disturbing the pellet. Then, pellets were resuspended in 2 ml shearing buffer and sonicated using a Focused Ultrasonicator M220 (Covaris). Sonication was performed in two rounds using two different sonication conditions: mild (peak power: 75, duty factor: 10.0, cycles per burst: 200, average power: 10.0, temperature range: 5-7°C) and intense (peak power: 75, duty factor: 25.4, cycles per burst: 200, average power: 19.1, temperature range: 5-7°C). Between sonication rounds, samples were centrifuged at 1,500 x g for 5 minutes. After sonication, cellular debris was precipitated by centrifugation at 14,000 x g for 20 minutes. An aliquot of clear supernatant was taken as ChIP input control (10 μg of chromatin). For chromatin immunoprecipiation, 25 μg of DNA were combined with 1 % Triton X-100, 150 mM NaCl and SPT4 antibody (1:100 dilution, Cell Signaling Technology, catalog number: 64828, lot number: 1). Samples were incubated rotating at 4°C overnight. Magnetic protein G Dynabeads (Invitrogen) were equilibrated by washing three times with IP buffer (1 % Triton, 0.1 % SDS, 1 mM EDTA, 10 mM Tris pH 8.0, 150 mM NaCl). The immunoprecipitation reactions were then incubated with the beads at 4°C for 90 minutes. The beads were subsequently washed twice with each TSE-150 (1 % Triton, 0.1 % SDS, 2 mM EDTA, 20 mM Tris pH 8.0, 150 mM NaCl) and TSE-500 buffer (1% Triton, 0.1% SDS, 2 mM EDTA, 20 mM Tris 8.0, 500 mM NaCl) and once with each LiCl (0.25 M LiCl, 1 % NP-40, 1 % sodium deoxycholate, 1 mM EDTA, 10 mM Tris pH 8.0) and TE buffer (1mM EDTA, 10 mM Tris pH 8.0). The beads were then incubated in 45 μl PK digestion buffer (20 mM Hepes pH 7.5, 1 mM EDTA pH 8.0, 0.5 % SDS) supplemented with 3 μl RNAse A (1 mg/ml stock, DNAse free, Thermo Fischer Scientific) at 37°C for 30 minutes. After addition of 5 μl proteinase K (1 mg/ml stock, Thermo Fischer Scientific), samples were incubated at 50°C for 30 minutes with periodic vortexing. Reverse crosslinking for both input control and ChIP samples was performed by adding NaCl to a final concentration of 0.3 M to the supernatant and incubating at 65°C overnight. DNA was purified using the Nucleospin Gel and PCR Clean up kit (Macherey-Nagel) by following the manufacturer’s instructions with slight modifications. After adding 5 volumes of buffer NTB, samples were centrifuged at 11,000 x g for 30 seconds and washed twice with 650 μl NT3 buffer. Buffer remnants were removed by centrifuging at 11,000 x g for 1 minute. DNA was eluted in adding 45 μl RNase-free water. Library preparation was performed as previously described (Tessarz *et al*, 2014). Libraries were sequenced on an Illumina Hi-Seq 2500 platform (paired-end, 50-bp reads).

### Cellular Fractionation

Cytoplasmic, soluble nuclear and chromatin-bound protein extracts were prepared by stepwise separation using Subcellular Protein Fractionation Kit for Tissues (Thermo Fisher Scientific, 87790). For this purpose, whole livers were harvested, snap-frozen in liquid nitrogen and stored at -80 °C. 200 mg of each homogenized liver sample were resuspended in 2 mL of ice-cold Cytoplasmic Extraction Buffer (CEB) supplemented with Halt Protease Inhibitor Cocktail (Thermo Fisher Scientific, 87786 at 1:100) and 1mM sodium metavanadate (NaVO_3_) and processed as outlined by the manufacturer’s protocol. Fractions were run on western blots and incubated with antibodies against RNA Pol II (Abcam, ab817), RNA Pol II Ser2 phospho (Abcam, ab5095) and RNA Pol II Ser 5 phospho (Abcam, ab5408).

### Tissue NET-seq (tNET-seq)

We modified the original NET-seq protocol (Mayer & Churchman, 2016) for application in murine liver tissue and named this new adaptation of the protocol tissue NET-seq (tNET-seq). The main change in the protocol pertains to the nuclei isolation. Fresh liver tissue were placed in ice-cold PBS, cut into smaller pieces and homogenize in 3 ml of nuclei isolation buffer (nuclei EZ lysis buffer (Sigma, NUC101), 1x Halt™ protease inhibitor cocktail (Thermo Fisher, 87786), 40 U RNasin (Promega, N2511), 25 μM α-amanitin (Sigma, A2263)) using a Dounce tissue homogenizer. In total, whole livers from 9 mice (3 per age group: young, middle-aged and aged) were processed immediately after sacrifice. The livers from young mice weighed between 1 and 1.5 g, while those from middle-aged and aged mice were heavier with 2-2.5 g. The entire procedure was performed on ice or at 4°C using RNase-free equipment. After complete homogenization, samples were transferred to a 15 ml tube. 3 ml of nuclei isolation buffer were added to the remaining pieces in the tissue homogenizer, homogenized further and transferred to same 15 ml tube. Following an incubation on ice for 5 min, samples were passed through a 70-μm cell strainer and nuclei were collected by centrifuging at 500 x g for 20 min. The nuclei pellet was resuspended in 4 ml of nuclei isolation buffer and incubated on ice for 5 min. Nuclei were collected by centrifuging at 500 x g for 5 min. To remove cytoplasmic remnants, the nuclei pellet was washed with 1600 μl nuclei wash buffer (1x PBS, 0.1 % (v/v) Triton X-100, 1 mM EDTA, 25 μM α-amanitin, 40 U RNasin, 1x Halt™ protease inhibitor cocktail) and centrifuged at 1,150 x g for 5 min. Washing was repeated with 800 μl nuclei wash buffer. Then, the pellet was gently resuspended in 200 µl of glycerol buffer (20 mM Tris-HCl (pH 8.0), 75 mM NaCl, 0.5 mM EDTA, 50 % (v/v) glycerol, 0.85 mM DTT, 25 μM α-amanitin, 10 U RNasin, 1x Halt™ protease inhibitor cocktail) using a cut 1,000-µl tip and transferred to a new 1.5-ml RNase-free microcentrifuge tube. Nuclei were lysed in 400 µl of nuclei lysis buffer 20 mM HEPES (pH 7.5), 300 mM NaCl, 1 % (v/v) NP-40, 1 M urea, 0.2 mM EDTA, 1 mM DTT, 25 μM α-amanitin, 10 U RNasin, 1x Halt™ protease inhibitor cocktail) and mixed by pulsed vortexing, followed by incubation on ice for 20 min. After assessing nuclei lysis of a small aliquot under a light microscope, samples were centrifuged at 18,500 x g for 2 min. The supernatant (nucleoplasmic fraction) was completely removed and the chromatin pellet was resuspended in 50 µl chromatin resuspension solution (PBS, supplemented with 25 μM α- amanitin, 20 U RNasin, 1x Halt™ protease inhibitor cocktail). A detailed protocol based on the Churchman lab protocol (Mayer & Churchman, 2016), with corresponding alterations for tissue is available from the corresponding author upon request.

### Computational analyses

#### Primary data processing

Primary sequencing data was quality controlled using FastQC (v 0.11.5) (Andrews & Others, 2010). Single-end tNET-seq reads were trimmed to 50-bp length using cutadapt (v 1.13) (Martin, 2011) and processed using custom Python scripts (adapted from https://github.com/BradnerLab/netseq). Paired-end ATAC-seq reads were trimmed to remove Tn5 transposase adapter sequence using cutadapt (v 1.13) (Martin, 2011) with the parameter –minimum-length=20. Reads were then aligned to the GRCm38 reference genome (Ensembl release 99) using Bowtie2 (v 2.4.1) (Langmead & Salzberg, 2012) by enabling soft clipping (--local) and the alignment of fragments up to 2 kb (-X 2000). Aligned reads were then filtered for high-quality (MAPQ > 10) and properly paired (samtools flag 0×2) (Li *et al*, 2009) reads. Finally, reads arising from PCR duplicates and those aligned to the mitochondrial genome were removed using PicardTools (v 2.21.4) (Institute, 2016) and samtools (v 1.10) in combination with grep, respectively.

ChIP-seq reads were aligned to the GRCm38 reference genome (Ensembl release 99) using Bowtie2 (v 2.4.1) by enabling soft clipping (--local). Aligned reads were then filtered for high quality (MAPQ > 10). Reads arising from PCR duplicates and those aligned to the mitochondrial genome were removed using PicardTools (v 2.21.4) and samtools (v 1.10) in combination with grep, respectively.

The quality of aligned reads was assessed using Rsamtools (v. 2.2.3) (Morgan, 2013), Additionally, fragment size distribution of ATAC-seq data was assessed using ATACseqQC (v. 1.14.4) (Ou *et al*, 2018).

#### Peak calling and annotation

ATAC-seq and ChIP-seq peaks were called on aligned and filtered BAM files using MACS2 (v. 2.2.7) (Gaspar, 2018). Where applicable, peak calling was performed in paired-end mode (-f BAMPE). For TF and histone ChIP-seq, the corresponding input libraries and total H3 ChIP-seq samples were used to determine the local background levels, respectively. Peaks displaying an FDR < 0.05 were considered as statistically significant.

The fraction of reads in peaks (FRiP) and fraction of reads in blacklisted regions (FRiBL) was determined using the R package ChIPQC (v. 1.21.0) (Carroll & Stark, 2017). Peaks falling in ENCODE blacklist regions were subsequently removed. Peaks were annotated using the R packages ChIPseeker (v. 1.26.2) (Yu *et al*, 2015) and ChIPpeakAnno (v 3.24.2) (Zhu *et al*, 2010). The promoter region was consistently defined as TSS +/-200 bp.

#### Gene selection

Annotation files for GRCm38 were retrieved from Ensembl (release 99). The genes included in the analysis were carefully selected to avoid contamination from transcription arising from other transcription units. Hence, only protein-coding genes were considered that are longer than 2 kb and non-overlapping within a region of 2.5 kb upstream of the TSS and downstream of the polyA site (n = 12,460). In case of multiple transcript isoforms, the most upstream annotated TSS and the most downstream annotated polyA sites were used. To ensure that only genes with sufficient coverage were included, the list was further filtered to contain only genes with RPKM > 1 (considering only uniquely aligned and filtered reads) (n = 3,280).

#### Differential accessibility analysis with ATAC-seq

Overlapping peaks that were called in different ATAC-seq samples were resolved by defining a consensus peak set containing non-redundant peaks present in at least two biological replicates regardless of condition (i.e. age). Reads overlapping consensus peak regions were recorded using featureCounts from the R package Rsubread (v. 2.0.1) (Liao *et al*, 2019). Differential accessibility analysis was performed using the R package DESeq2 (v. 1.26.0) (Love *et al*, 2014). Regions with an FDR < 5 % were considered to be statistically significant.

#### Differential expression analysis with RNA-seq

Differential expression analysis was performed using DESeq2 (v. 1.26.0) (Love *et al*, 2014). Genes with an FDR < 5 % were considered to be statistically significant.

#### Recording Pol II density at nucleotide resolution

Pol II density was calculated by recording the genomic position of the 5’ end of each tNET-seq read, which corresponds to the 3’ end of the original nascent RNA and represents the exact genomic position of Pol II. For this, bedtools (v. 2.29.2) (Quinlan & Hall, 2010) genomecov with the parameters -dz and -5 was used.

#### Differential transcription analysis with tNET-seq

Pol II density in gene bodies of tNET genes was retrieved using bedtools (v. 2.29.2) (Quinlan & Hall, 2010) intersect and the read count matrix of genome-wide Pol II densities. For this, the gene body was defined as 200 bp downstream of the TSS to 200 bp upstream of the TES. To identify differentially transcribed genes, differential analysis of Pol II density in gene bodies was performed using the R package DESeq2 (v. 1.26.0) (Love *et al*, 2014). Here, only sense transcription was considered (i.e. tNET-seq reads sharing the same orientation as the annotation). Genes with an FDR < 5 % were considered to be statistically significant.

#### Trajectory analysis of nascent transcription with tNET-seq

To estimate nascent transcription trajectories during aging, we performed a likelihood-ratio test using the R package DESeq2 (v. 1.26.0) (Love *et al*, 2014). Regions with an FDR < 5 % were considered to be statistically significant. Then, we identified common patterns using the R package DEGreport (v. 1.26.0) (Pantano & Others, 2019). In brief, rlog-transformed counts of significantly differentially transcribed genes were used for calculating pair-wise gene expression among conditions using Kendall’s rank. Divisive hierarchical clustering (DIANA) was then used on the gene-gene distance matrix for identifying groups of genes with similar trajectories. Z-scores of these genes were visualized.

#### Pol II pausing index

To quantify promoter-proximal pausing, we calculated the Pol II pausing index, which is defined as the ratio between the average Pol II density in the promoter-proximal region (defined here as TSS +/- 200 bp) over that in the gene body (defined here as TSS + 200 bp to TES - 200 bp).

For computing the Pol II pausing index, RPM-normalized Pol II coverage files were generated using bedtools (v. 2.29.2) (Quinlan & Hall, 2010) genomecov with the parameters -dz, -5 and -scale 1/number of aligned reads in mio. The number of aligned reads after pre-processing was obtained using samtools view (v 1.10) (Li *et al*, 2009) with the flag -c and -F 260 to output the number of primary aligned reads only.

The RPM-normalized count matrices were intersected with annotation files for both promoter-proximal and gene body regions using bedtools (v. 2.29.2) (Quinlan & Hall, 2010) intersect. After normalizing for region length, the pausing index was calculated by dividing the mean normalized Pol II density in the promoter-proximal region by the mean normalized Pol II density in the gene body region for each tNET gene. Here, only sense transcription was considered (i.e. tNET-seq reads sharing the same orientation as the annotation). Extreme data points were removed from the pausing analysis by retaining only genes with a pausing index ≤ 10 in all samples (n = 109 genes removed).

#### Pol II pausing index and promoter accessibility

To allow for a direct comparison between promoter accessibility and the Pol II pausing index, the ATAC-seq data was processed analogous to the tNET-seq data. In brief, RPM-normalized chromatin accessibility was calculated by recording the genomic position of the 5’ end of each ATAC-seq read using bedtools (v. 2.29.2) (Quinlan & Hall, 2010) genomecov as described above. We then computed chromatin accessibility in promoter-proximal regions (TSS +/- 200 bp) using bedtools (v. 2.29.2) intersect. After normalizing for region length, the log_2_ fold change in promoter accessibility in aged versus young mice was compared to the log_2_ fold change in pausing index in aged versus young mice.

#### Identification of active enhancers in murine liver tissue

For identifying active enhancers in murine liver tissue, we combined our accessibility data (ATAC-seq) with publicly available histone modification data (H3K27ac and H3K4me3 ChIP-seq). For each dataset, we defined a consensus peak set containing peaks present in all samples regardless of biological condition (i.e. age). Enhancers were then identified as H3K27ac peaks that (i) do not overlap H3K4me3 peaks, (ii) do not fall within TSS +/-1 kb, and (iii) overlap accessible sites identified through ATAC-seq. This resulted in the identification of 8,855 enhancer regions active in murine liver tissue.

#### Functional enrichment analysis

GO databases were queried using the R package clusterProfiler (v. 3.14.3) (Yu, 2018). To test for over-representation, the complete gene list of the mouse database (Mm.eg.db, v. 3.12.0) served as background. After removing semantic redundancy, GO terms were ranked by adjusted p-value (Benjamini-Hochberg procedure, FDR < 0.05).

#### Visualization

For visualization purposes, aligned and filtered BAM files were converted to bigwig coverage tracks using Deeptools (v. 3.5.1) (Ramírez *et al*, 2016). For ATAC-seq and tNET-seq, a bin size of 1 bp was used, while ChIP-seq reads were extended and counted in 10-bp bins. Bigwig files were normalized to 1x coverage (--normalizeUsing RPGC). For tNET-seq, only the position of the 5’ end of the sequencing read was recorded (--Offset 1).

Metagene profiles and heatmaps of mean signal enrichment were generated using Deeptools (v 3.5.1) and files normalized to 1x coverage as input. For tNET-seq, reads sharing the same or opposite orientation with the annotation were assigned as sense or antisense, respectively. Biological replicates were visualized either separately or merged prior to visualization. Sample-by-sample correlation and principal component analyses were performed using rlog-transformed read counts via DESeq2 (v. 1.26.0) (Love *et al*, 2014). Single-gene sequencing tracks were visualized with Integrative Genomics Viewer (v. 2.8.0) (Robinson *et al*, 2011).

#### Reproducibility and statistical analysis

R v. 3.6.3 and Python v. 3.9.0 were used for all analyses. Statistical parameters and significance are reported in the figures and figure legends. Whenever possible, we used non-parametric statistical tests to avoid assuming normality of data distributions.

## DATA AVAILABILITY

All data generated in this study is available at GEO, accession number: XXX. Tabula Muris senis bulk RNA-seq data (Schaum *et al*, 2020) was retrieved from Figshare as raw count table (https://figshare.com/projects/The_murine_transcriptome_reveals_global_aging_nodes_with_organ-specific_phase_and_amplitude/65126). Additionally, we analyzed publicly available histone ChIP-seq data for H3, H3K27me3 and H3K4me3. Raw data originating liver tissue of 3, 12 and 29-month-old male mice was retrieved from the NCBI BioProject database (PRJNA281127) (Benayoun *et al*, 2019).

## AUTHOR CONTRIBUTIONS

Conceptualization: M.B. and P.T.; Methodology: M.B., D.G., C.N.; Investigation: M.B., D.G., C.N.; Formal Analysis: M.B.; Supervision: P.T.; Funding Acquisition: P.T.; Project Administration: P.T.; Writing of Manuscript: M.B., P.T., with input from all authors

## ACKNOWLEDGEMENTS

We would like to thank all members of the Tessarz lab for continuous discussion. We are grateful to A. Pouikli for critical reading of the manuscript. We thank the following core facilities for expert technical assistance: Comparative Biology of the MPI for Biology of Ageing, Cologne for mouse husbandry and the Genomic Core Facility at the MPI for Molecular Genetics, Berlin for sequencing. This work was funded by the Max Planck Society (to P.T.) and the Deutsche Forschungsgemeinschaft (DFG, under Germany’s Excellence Strategy – EXC 2030 – 390661388) (to P.T.). M.B. received support by the Cologne Graduate School of Ageing Research. Models were drawn using Biorender.

## SUPPLEMENTARY FIGURES

**Supplementary Figure 1:**
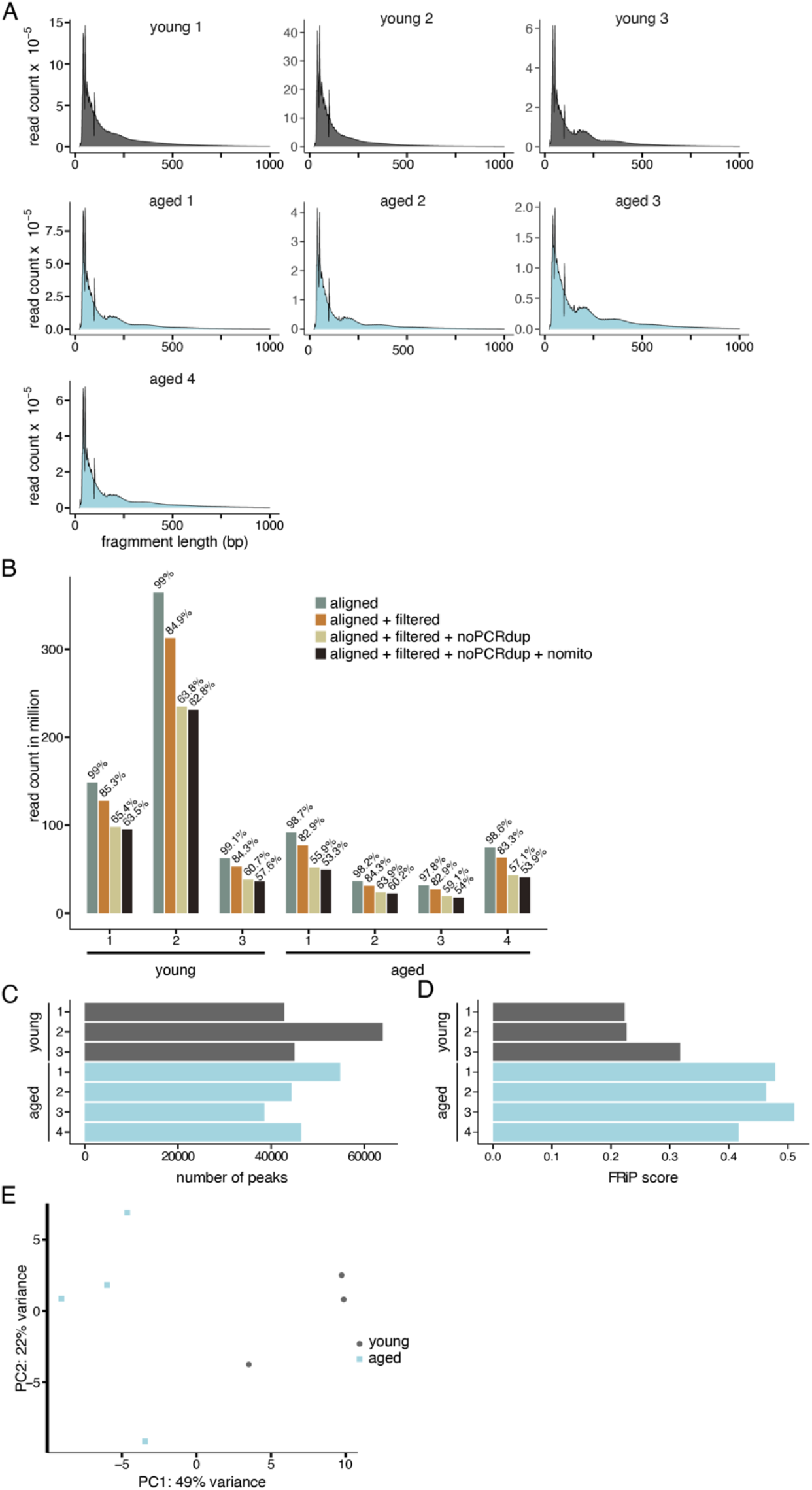
A. Fragment size distribution of reads passing filtering criteria for each ATAC-seq library. Each curve represents one biological replicate (n = 3 and 4 young and aged mice, respectively). B. Number of ATAC-seq reads successfully aligned and passing filtering criteria. Aligned reads were filtered for high quality (MAPQ > 10) and PCR duplicates and mitochondrial reads were removed. C. Number of peak regions identified in each ATAC-seq library. D. Signal-to-noise ratio assessed by the fraction of reads in peaks (FRiP). E. Principal component analysis of chromatin accessibility profiles. Rlog-normalized read counts (DESeq2) in consensus peak regions identified by ATAC-seq are depicted. Percentage of variance accounted for by each principal component is indicated.

**Supplementary Figure 2:**
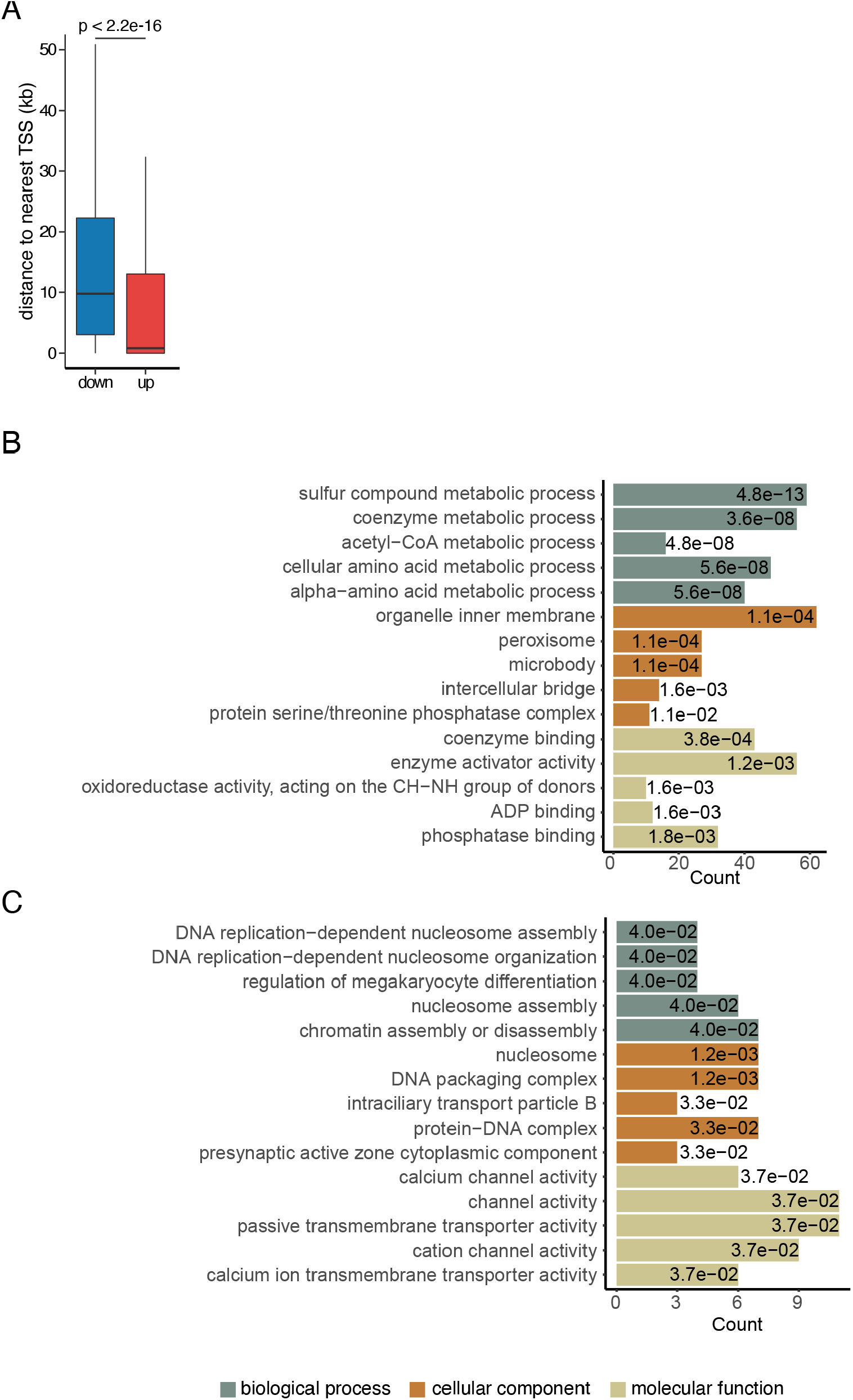
A. Distance of differentially accessible sites to the nearest annotated TSS. P-value was calculated using a two-sided Wilcoxon rank-sum test. B, C. GO term enrichment analysis for genes with increased (B) and decreased (C) promoter accessibility in the liver of aged mice. Only genes with a differentially accessible TSS were included in the analysis (n = 1,945 genes in total; 1,704 with increased and 241 with decreased promoter accessibility). Only the top five enriched GO terms ranked by adjusted p-value (Benjamini-Hochberg (BH) procedure, FDR < 0.05) are displayed. BH-adjusted p-values are reported for each GO term inside the respective bar.

**Supplementary Figure 3:**
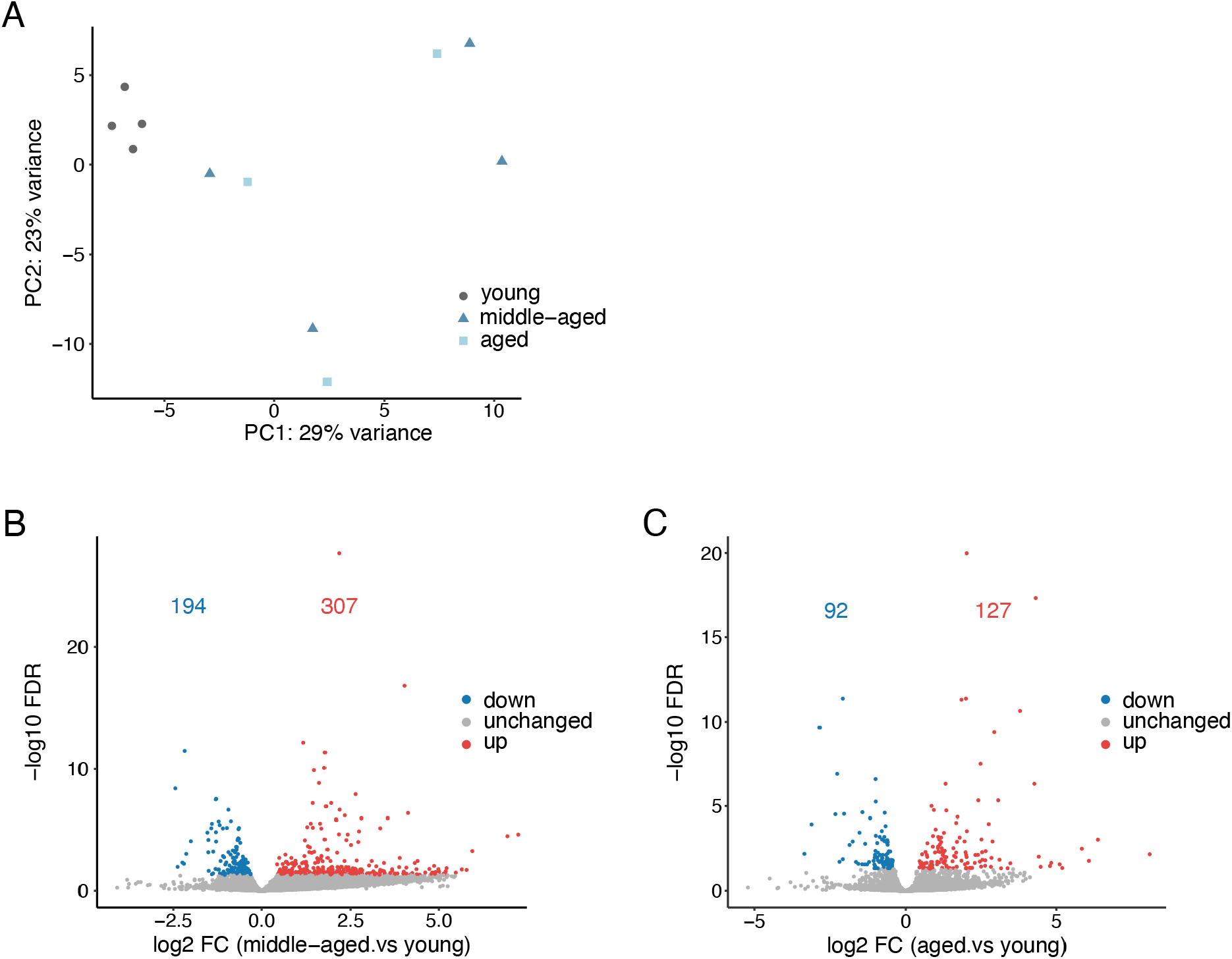
A. PCA scatter plot of steady-state gene expression profiles assessed by RNA-seq. Percentage of variance accounted for by each principal component is indicated. B. Volcano plot of differentially expressed genes comparing middle-aged relative to young animals (FDR < 0.05, Wald test). 307 genes were up-regulated (red) and 194 down-regulated (blue). C. Volcano plot of differentially expressed genes in the liver of aged versus young animals (FDR < 0.05, Wald test). 127 genes were up-regulated (red) and 92 down-regulated (blue).

**Supplementary Figure 4.**
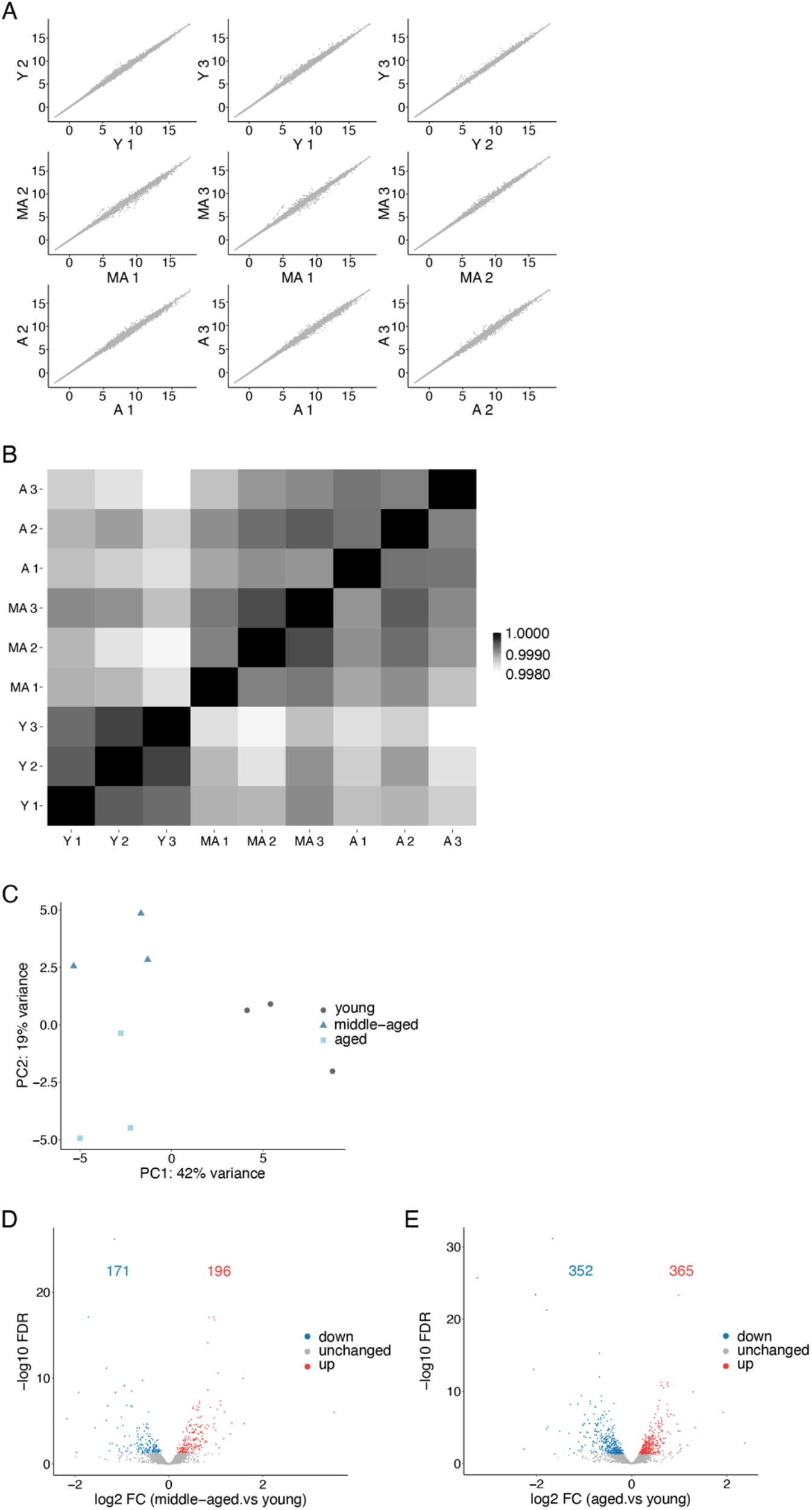
A. Pairwise comparison of tNET-seq samples. Normalized read counts (rlog transformation, DESeq2) in non-overlapping, protein-coding genes above 2 kb in size are reported. B. Heatmap of replicate correlations using normalized read counts as in A. The color code represents Pearson correlation coefficient. C. Principal component analysis of nascent transcription assessed by tNET-seq. PCA was performed with the normalized read counts (rlog transformation, DESeq2) in gene bodies of non-overlapping, protein-coding genes above 2 kb in size. The percentage of variance accounted for by each principal component is indicated. D. Volcano plot of differentially transcribed genes (gene-body Pol II density) in middle-aged relative to young animals (FDR < 0.05, Wald test). 196 genes were up-regulated (red) and 171 down-regulated (blue). E. Volcano plot of differentially transcribed genes (gene-body Pol II density) in aged relative to young animals (FDR < 0.05, Wald test). 365 genes were up-regulated (red) and 352 down-regulated (blue). Y, young; MA, middle-aged; A, aged.

**Supplementary Figure 5:**
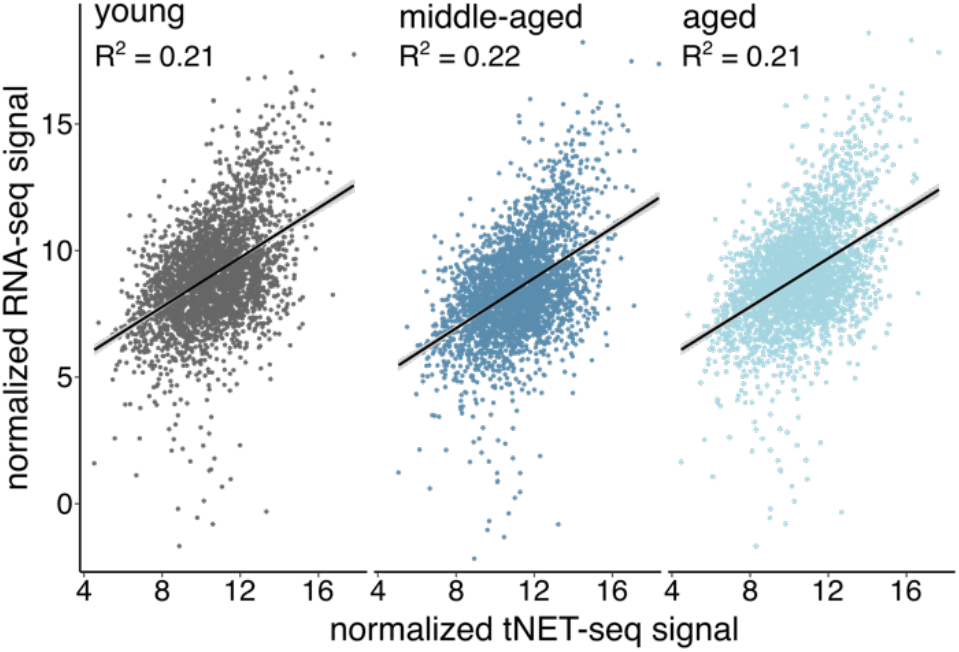
Correlation between nascent (gene-body Pol II density, tNET-seq) and steady-state transcription (RNA-seq). Mean normalized read counts (rlog transformation, DESeq2) of merged biological replicates from young, middle-aged and aged animals are reported.

**Supplementary Figure 6:**
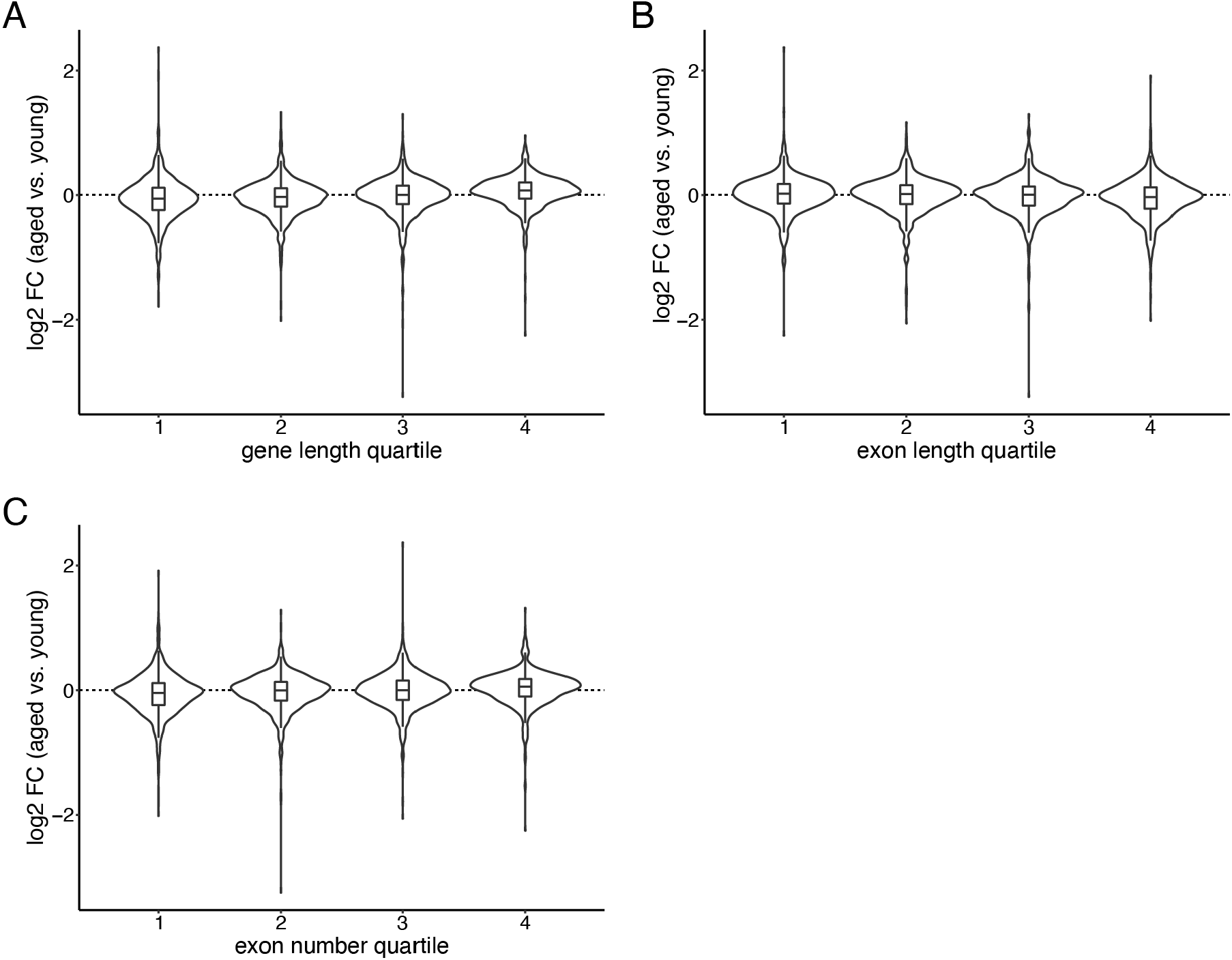
Changes in nascent transcription (gene-body Pol II density, tNET-seq) in aged versus young animals in relationship to gene length (A), median exon length (B) or exon number per gene (C). The gene length was calculated as the total exonic length after reducing a gene’s exons to a non-overlapping set. Number of genes per quartile: 819.

**Supplementary Figure 7:**
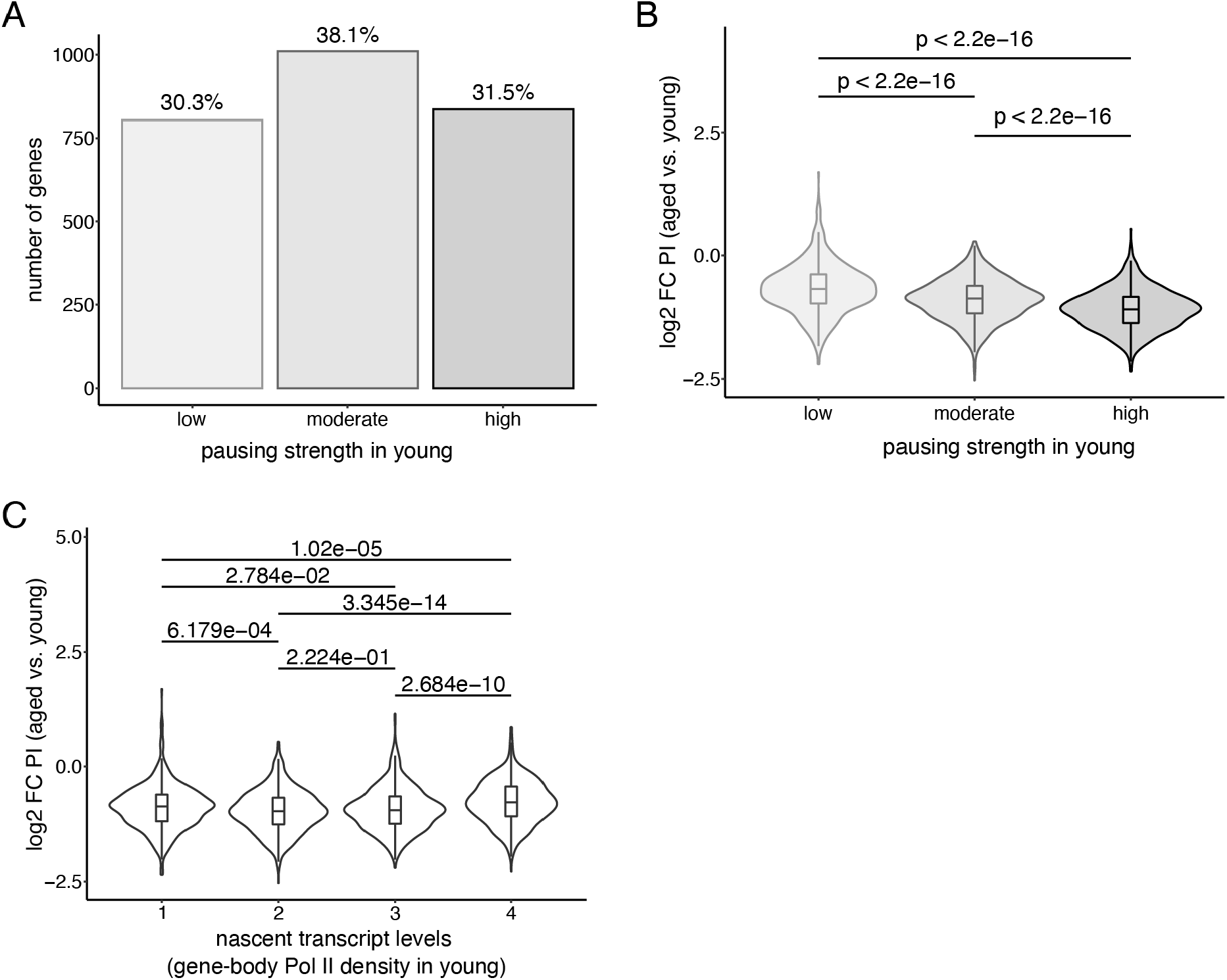
A. Bar plot of PIs quantifying the extent of promoter-proximal pausing in young mice. The 2,650 genes were divided into three groups based on the PI in young animals: highly paused (PI ≥3; n = 836), moderately paused (1.5 ≤ PI < 3; n = 1,010), and lowly-paused (PI < 1.5; n = 804). B. Violin and box plot of change in PI of aged versus young liver. Genes were grouped by PI as in A. C. Violin and box plots of PI change in aged versus young animals. Genes were divided into four equal-sized groups based on their level of nascent transcription (gene-body Pol II density) in the liver of young mice. p-values were calculated using a two-sided Wilcoxon rank-sum test.

**Supplementary Figure 8:**
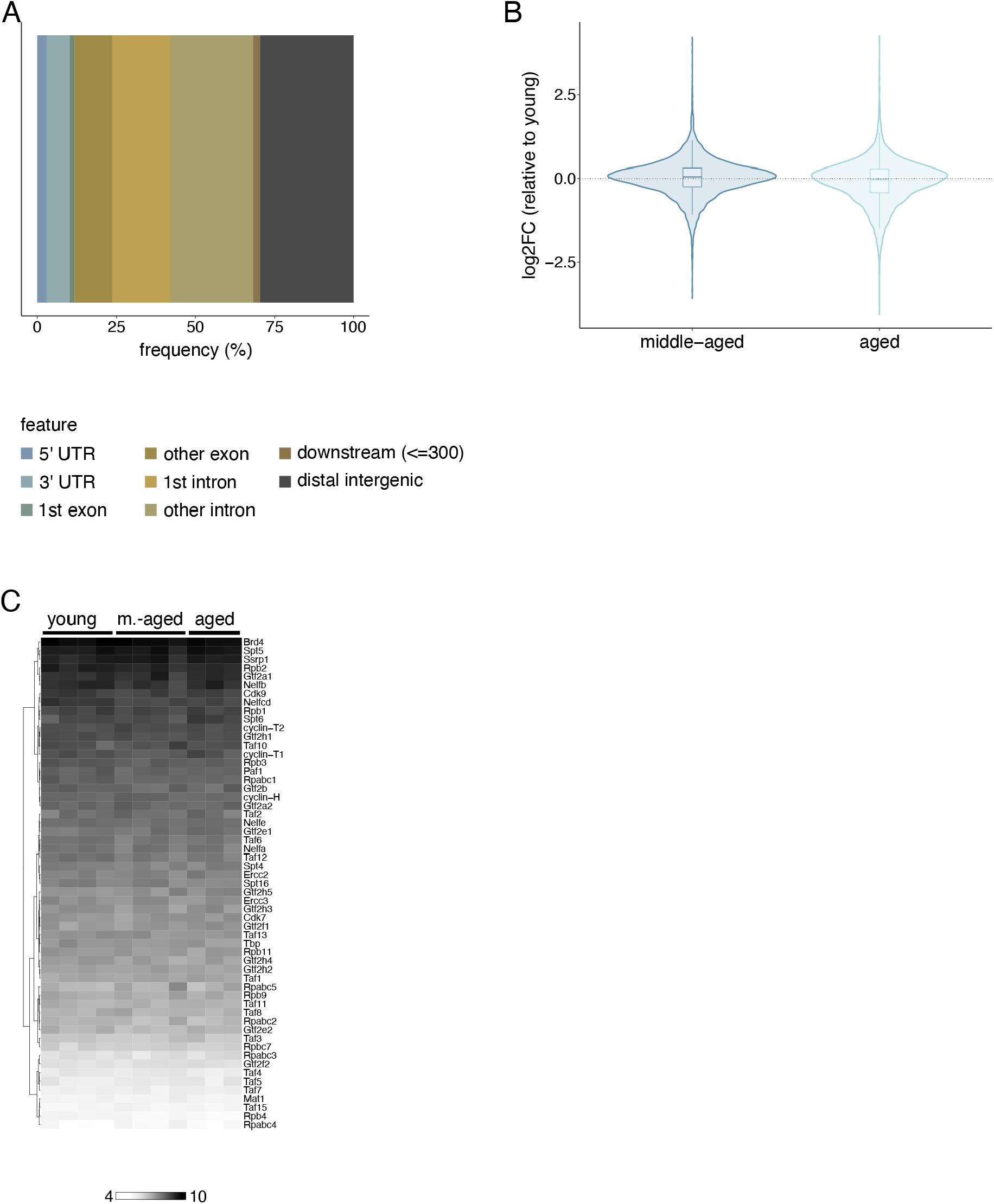
A. Genomic distribution of identified active enhancers. B. Violin and boxplots of log2-fold changes in Pol II density at enhancer regions of middle-aged and aged animals relative to young ones. C Heatmap of steady-state mRNA levels (RNA-seq) of relevant transcription regulators in young, middle-aged and aged mice. Normalized read counts (rlog transformation, DESeq2) are reported.

